# Enhancers require an active resetting phase after transcriptional activation

**DOI:** 10.64898/2026.09.19.752844

**Authors:** Arif Hussain Najar, Rajat Mann, Subhashis Indra, Deepanshu Soota, Paramita Kundu, Sundarraj Nidharshan, Rakesh Netha Vadnala, Dimple Notani

**Affiliations:** National Centre for Biological Sciences (NCBS), Bengaluru, India; The University of Trans-Disciplinary Health Sciences and Technology, IVRI Road, Bengaluru, India; School of Biotechnology, Amrita Vishwa Vidyapeetham, Kollam, India

## Abstract

Signal-responsive enhancers must activate transcription and then return to a competent but inactive state, yet whether this transition is passive or actively driven has remained unresolved. We define this process as enhancer resetting and identify the androgen receptor (AR) as the factor that executes it as an endogenous response to estrogen signaling, independent of exogenous androgen. As estrogen receptor-α (ERα) undergoes ligand-induced proteasomal degradation during late-phase signaling, AR progressively accumulates in the nucleus and preferentially occupies highly active, persistent ERα enhancers in a transcriptionally inactive manner. This late-phase AR binding evicts the pioneer factor FOXA1 from chromatin while preserving baseline accessibility, a handoff mechanism that decouples chromatin openness from active transcription. Conversely, augmenting AR activity by DHT treatment or AR overexpression accelerates FOXA1 eviction, while disrupting AR chromatin binding, either by AR knockdown or a DNA-binding-deficient mutant, prevents it, leaving enhancers in an aberrantly permissive state that drives amplified ERα rebinding and transcriptional hyperactivation upon subsequent estrogen stimulation. These findings establish active enhancer resetting as a mechanism safeguarding the fidelity of repeated transcriptional responses. Failure of this mechanism may underlie the transcriptional dysregulation that drives tumor progression and endocrine therapy resistance in ERα-positive breast cancer, where estrogen signaling is chronic and cyclic.

## Introduction

Signal-dependent transcription factors drive gene expression in precise temporal pulses, orchestrating rapid cellular adaptations to developmental, physiological, and environmental cues. Central to these responses are enhancers, which integrate extracellular signals by recruiting signal-dependent transcription factors, coactivators, and the transcriptional machinery. Considerable progress has been made in defining how enhancers are activated, converging on a hierarchical model in which pioneer transcription factors establish accessible chromatin and thereby permit inducible transcription factors to engage and drive transcription (Carroll et al. 2005, Lupien et al. 2008, Zaret and Carroll 2011). Yet many signaling pathways operate in repeated cycles; enhancers must not only activate transcription efficiently but also be restored to a transcriptionally competent state that permits faithful responses to subsequent stimulation. How enhancers are reset to this transcriptionally competent state following activation remains largely unknown.

Enhancer resetting is conceptually distinct from enhancer poising. Poised enhancers (marked by H3K27me3 and H3K4me1), studied primarily in developmental contexts, exist in an inactive, primed state prior to any activation and are defined by chromatin signatures that anticipate future lineage-specific gene expression (Creyghton et al. 2010, Rada-Iglesias et al. 2011). Signal-responsive enhancers, by contrast, repeatedly cycle between active and inactive states throughout the lifetime of a differentiated cell (Fukaya et al. 2016). Following each signaling cycle, they must dismantle the active transcriptional apparatus without permanently losing regulatory competence. Failure to reset the enhancer state would be expected to either promote persistent transcriptional memory and inappropriate hyperactivation or impair future responsiveness altogether (Tan et al. 2018, Yang et al. 2020). Whether this transition is a passive consequence of transcription factor dissociation or an actively regulated process has not been resolved. We define enhancer resetting as the active restoration of an enhancer from a transcriptionally active state to a transcriptionally competent but inactive one, enabling faithful responses to subsequent rounds of signaling.

Estrogen signaling provides an ideal model to investigate enhancer resetting because it generates one of the most dynamic enhancer-driven transcriptional responses. Upon estradiol (E2) stimulation, estrogen receptor-α (ERα) rapidly occupies thousands of distal enhancer regions genome-wide (Li et al. 2013). There, it recruits Mediator, p300, and RNA polymerase II to drive transcriptional programs governing mammary epithelial identity, proliferation, and differentiation. Both genomic occupancy and target gene transcription peak within the first hour of stimulation (Hah et al. 2011, Liu et al. 2014, Saravanan et al. 2020). This acute transcriptional surge is short-lived: by three hours post-stimulation, ERα undergoes ligand-induced proteasomal degradation (Nawaz et al. 1999), resulting in a widespread loss of genomic ERα occupancy and a return of target gene expression to baseline levels by 24h despite continued hormone exposure (Dzida et al. 2017).

ERα recruitment during this early phase (peak phase) is critically dependent on the pioneer factor FOXA1, which establishes accessible chromatin and licenses receptor binding (Cirillo et al. 2002, Carroll et al. 2005, 2006, Hurtado et al. 2011). FOXA1 is conventionally regarded as a stable, lineage-defining feature of the ERα cistrome rather than one subject to dynamic regulation (Lupien et al. 2008). While the mechanisms governing acute ERα activation and subsequent clearance are well delineated, how cells reset these transiently hyperactive genomic regions between signaling cycles, and whether FOXA1 occupancy itself remains stable or is dynamically remodeled during this transition (Swinstead et al. 2016, Glont et al. 2019) is a fundamental, unresolved question in hormone biology. The one with direct bearing on how transcriptional memory is prevented from accumulating unchecked across repeated signaling cycles.

The Androgen Receptor (AR), a lineage-defining nuclear receptor expressed alongside ERα in most estrogen-responsive breast tumors, is conventionally recognized as a tumor suppressor in this context (Peters et al. 2009, Hickey et al. 2021). In the presence of canonical androgenic ligands, AR directly competes with ERα and redirects coactivators away from estrogen-responsive elements, thereby antagonizing active ERα-driven transcription (Hickey et al. 2021). Notably, AR and FOXA1 have been shown to engage in a similarly complex, competitive relationship in prostate cancer (Sahu et al. 2011, Jin et al. 2014), raising the possibility that analogous AR-FOXA1 dynamics may operate in other hormone-responsive tissues. This model, however, rests on androgen-replete conditions that do not reflect the hormonal environment of most estrogen-responsive breast tumor models, in which intracellular steroidogenesis favors the conversion of androgens to estrogens rather than the reverse (Payne and Hales 2004). Notably, AR accumulates progressively during prolonged estrogen signaling even in the complete absence of androgen stimulation (D’Amato et al. 2016), raising the possibility that AR serves functions beyond its canonical role as a ligand-activated transcription factor, specifically, across the distinct temporal phases of estrogen signaling, and particularly during the late transition phase following ERα degradation, a phase that existing models of AR-ERα antagonism have not addressed.

Here, we identify a chronic role for AR as an essential enhancer-resetting factor during late-phase estrogen signaling. Across an extended timeline of E2 exposure (0-24 h), while ERα levels drop, AR progressively accumulates in the nucleus and preferentially occupies the subset of highly active ERα regulatory elements, which we have defined as “persistent enhancers” (Saravanan et al. 2020). Unexpectedly, AR binding at the late phase (24 h) is transcriptionally silent, yet is required to clear the pioneer factor FOXA1 from chromatin while preserving baseline chromatin accessibility, a “handoff” mechanism that decouples accessibility from active transcription. By disrupting AR binding through siRNA knockdown or a DNA-binding-deficient mutant (AR-DBM), we demonstrate that failure to evict FOXA1 leaves enhancers in a hyper-competent state, resulting in amplified ERα rebinding and transcriptional hyperactivation upon subsequent estrogen exposure. Together, our findings establish active enhancer resetting as a fundamental mechanism safeguarding the fidelity of repeated transcriptional responses, and identify AR as an enhancer-resetting factor that resets strong regulatory elements and restrains hyperactive estrogen signaling across successive signaling cycles.

## Results

### AR accumulates in the nucleus and occupies chromatin at the late phase of estrogen signaling

Estrogen elicits a rapid and transient increase in transcriptional activity, with gene expression upregulated as early as 10 minutes following E2 addition (Hah et al. 2011, Saravanan et al. 2020). The transcriptional dynamics of most E2-regulated genes closely mirror ERα binding patterns at regulatory sites, with both reaching peak activity within 1 h and subsequently declining from 3 h of signaling onward (Hah et al. 2011). To investigate the E2 program during the transition from the early to the late phase of signaling, we utilized published ChIP-seq data for ERα at different time points of E2 treatment (0-∼21h) (Dzida et al. 2017). As reported earlier, ERα binding to chromatin peaked at 40 minutes following ligand stimulation, declined to near-basal levels by 3 h, and returned to basal levels by 1280 minutes (∼21 h) (Fig. 1A). Consistent with this, the transcription of ERα-associated genes also mirrored the ERα binding, peaking at 40 minutes and decreasing by 160 minutes (3h) (Fig. 1B). The loss of ERα binding across the genome after 3 h has been attributed to proteasomal degradation of ERα (Nawaz et al. 1999, Reid et al. 2003, Fan et al. 2004). Similar to these observations, analysis of whole-cell extracts revealed a progressive reduction in ERα levels at 1, 3, and 24 h following estrogen stimulation, with ERα levels becoming extremely low by 24 h (Fig. 1C, upper panel).

**Fig. 1:**
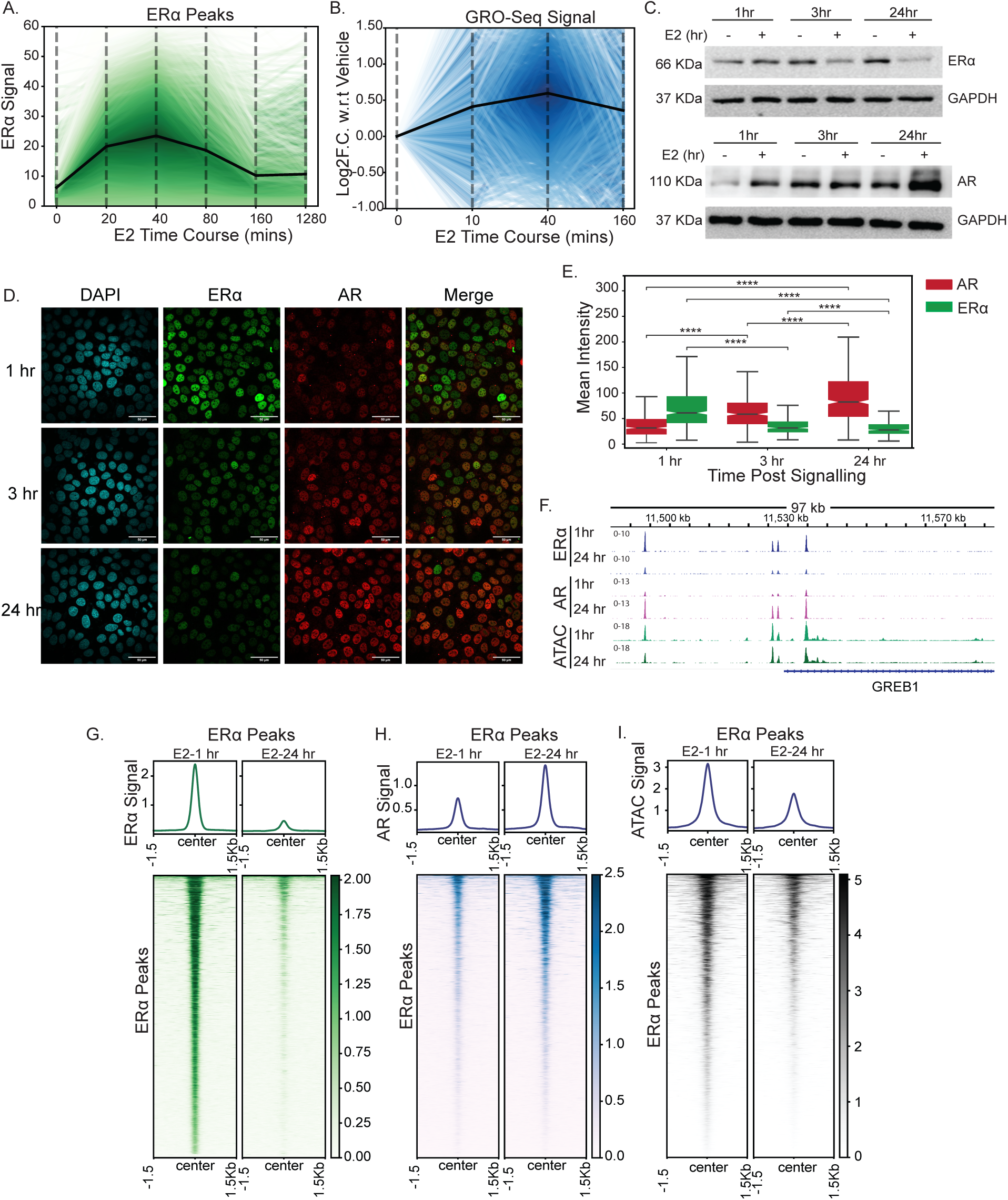
AR accumulates in the nucleus and occupies chromatin at the late phase of estrogen signaling: **A.** ChIP-seq signal of ERα peaks at 0, 20, 40, 80, 160, and 1280 min post-E2 signaling. The x-axis denotes the different time points. y-axis denotes ChIP-seq normalized counts. Each line depicts an individual peak. The black line indicates the mean curve. The hue of the line denotes the z-score for the ChIP-seq signal at 40min. **B.** GRO-seq signal of genes associated with ERα peaks at 0, 10, 40 and 160 min post E2 signaling. The x-axis denotes the different time points post signaling. The y-axis denotes log2FC. w.r.t 0 min. Each line depicts an individual gene. The black line indicates the mean curve. The hue of the line indicates the z-score for log2FC. values at 40min. **C.** Western blot of MCF7 cell lysate showing ERα (Upper panel) and AR levels (lower panel) at 1, 3, and 24 h in the presence (+) and absence (-) of E2. GAPDH levels are used as a loading control. **D.** Confocal images of MCF7 cells at 1, 3, and 24 h of E2 signaling immunostained for ERα and AR. The scale bar represents 10 µm. DAPI is used for nuclear staining. **E.** Plot of mean nuclear intensity of ERα and AR at 1, 3, and 24 h of E2 signaling. The boxplots depict the minimum (Q1-1.5IQR), first quartile, median, third quartile, and maximum (Q3 + 1.5IQR) without outliers. p-values are calculated using the two-tailed Mann-Whitney-Wilcoxon test. (ns P > 0.05; *P < 0.05; **P < 0.01; ***P < 0.001; ****P < 0.0001). **F.** IgV snapshot of ERα, AR and ATAC-seq signal at 1 and 24 h of E2 signaling on the GREB1 gene locus. **G.** Heatmap showing the ERα signal at 1 and 24 h of E2 signaling on ERα peaks. **H.** Heatmap showing the AR signal at 1 and 24 h of E2 signaling on ERα peaks. **I.** Heatmap showing the ATAC-seq counts at 1 and 24 h of E2 signaling on ERα peaks.

The androgen receptor (AR) is a lineage-defining nuclear receptor that is co-expressed with ERα in the majority of estrogen-responsive breast tumors and has traditionally been considered to exert tumor-suppressive functions (Peters et al. 2009, Hickey et al. 2021). Upon binding canonical androgenic ligands, AR can antagonize ERα activity by redirecting liganded AR and shared coactivators toward AR target genes, thereby attenuating ERα-driven transcription (Hickey et al. 2021). However, breast tumors are exposed to sustained estrogenic signaling, and how AR is regulated during the temporal decline in ERα activity remains unclear. We therefore asked whether AR levels change as ERα transcriptional activity declines at later phase of estrogen signaling. To address this, we examined AR protein levels across the estrogen stimulation time, i.e. 1, 3, and 24 h after E2 treatment. Surprisingly, in contrast to ERα, AR protein levels increased from 1 to 24 h (Fig. 1C lower panel). At 24 h, immunofluorescence revealed a reciprocal change in nuclear receptor abundance, with increased nuclear accumulation of AR concomitant with reduced nuclear ERα (Fig. 1D-E).

To assess whether nuclear-accumulated AR indeed binds to chromatin during the late phases of signaling, we performed ChIP-seq for both ERα and AR at 1 and 24 h after E2. As previously shown, we observed a decrease in both the level and the number of ERα binding sites in the genome (Saravanan et al. 2020). The number of ERα binding sites decreased markedly from 3,088 at 1 h to 288 at 24 h following E2 treatment. We identified 2,826 ERα peaks lost (91.5%), 262 peaks shared between the two time points, and only 26 peaks gained at 24 h. (Fig. 1F-G; Supp Fig. 1A-B). Strikingly, AR binding increased markedly at 24 h, rising from 2,808 binding sites at 1 h to 4,954 sites at 24 h (76% increase) closely mirroring the increase in AR protein levels. Of these, 1,738 sites were shared between the two time points (Supp Fig. 1C-D). Since, our experimental set up only contained estrogen not testosterone, accumulated AR and its increased occupancy at chromatin was intriguing. Given the inverse relationship between AR and ERα levels, we asked whether AR is enriched at ERα-bound regions. Indeed, AR binding at ERα peaks was low during the early phase of signaling (1 h) but increased substantially during the late phase (24 h), coincident with a decrease in ERα binding from 1 to 24 h (Fig. 1F-H). Since both TFs showed opposing binding dynamics at the same ERα-bound sites, we asked whether chromatin accessibility also changed during the switch from ERα to AR binding. ATAC-seq profiling revealed a reduction in chromatin accessibility during the late phase; however, despite the decrease, the majority of ERα-bound sites remained accessible even as ERα binding was substantially reduced (Fig. 1I).

### AR occupies active ERα regulatory elements at the late phase of signaling

Peak calling of the AR datasets revealed that approximately 39% (1,198 of 3088) of the ERα-bound regions at 1 h were subsequently occupied by AR at 24 h (Fig. 2A). Thus, while ERα binding was lost during the late phase, a subset of ERα-bound regions gained AR binding (Fig. 2C), whereas the remaining regions lost ERα binding without acquiring AR (Fig. 2B). While AR also bound to non-ERα sites at 24 h, AR binding intensity was markedly higher at AR-ERα overlapping regions than at AR-specific sites (Fig. 2D). We therefore compared these two classes of ERα-bound regions, those that gained AR binding at 24 h (ERα-AR overlapping regions) and those that did not (non-overlapping), to identify features associated with the transition from ERα to AR binding.

**Fig. 2:**
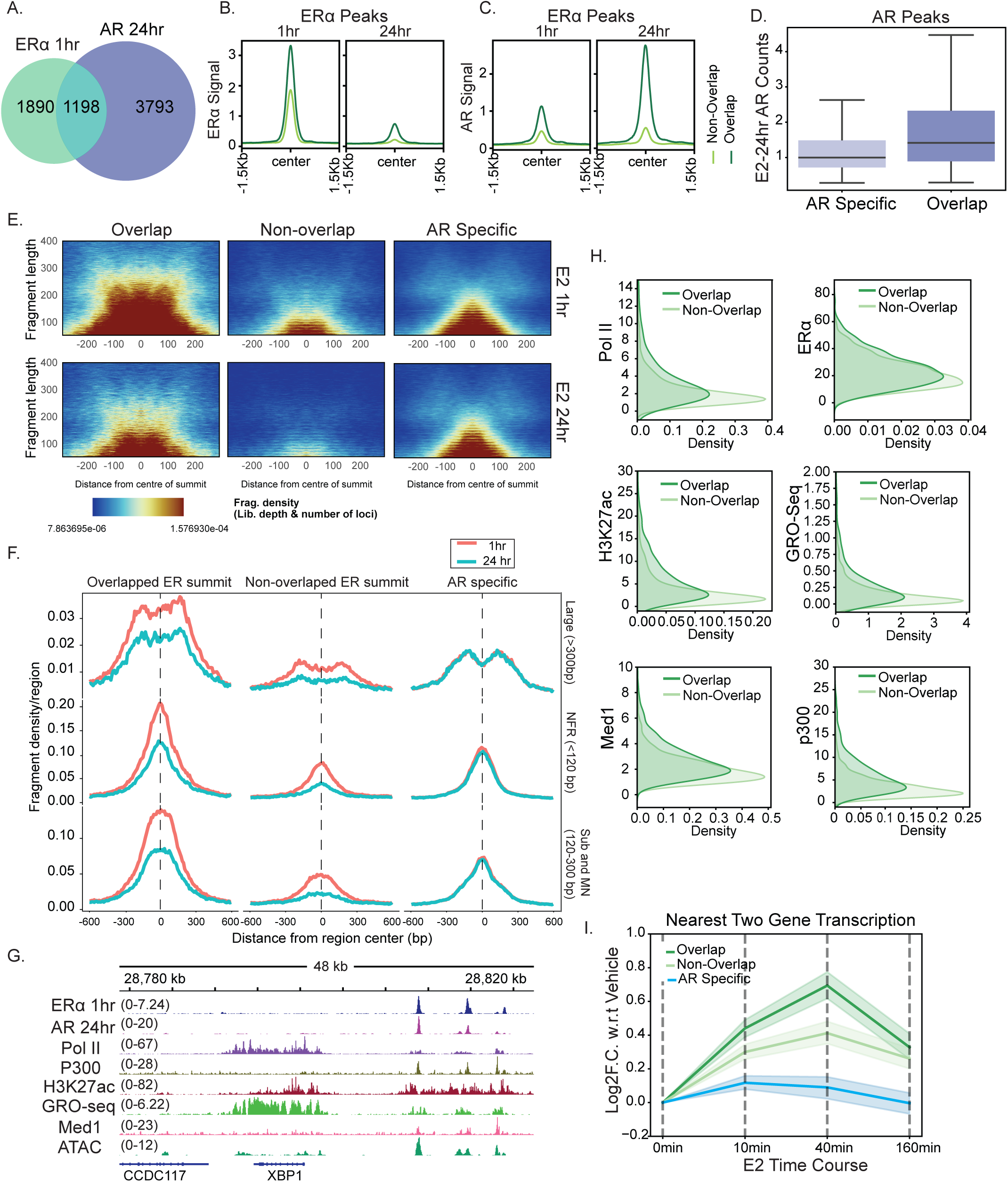
AR occupies active ERα regulatory enhancers at the late phase of signaling. **A.** Venn diagram showing overlap of ERα peaks at 1h and AR peaks at 24hr. **B.** Profile plot showing binding of ERα on ERα peaks at 1 and 24 h post-signaling. The dark green curve indicates Overlapping sites and light green shows non-overlapping sites. **C.** Profile plot showing binding of AR on ERα peaks at 1 and 24 h post-signaling. The dark green curve indicates Overlapping sites and light green shows Non-overlapping sites. **D.** Boxplot of AR signal at 24 h of E2, on AR: ERα Overlapping peaks and Peaks specific to AR at 24 h. The boxplots depict the minimum (Q1-1.5IQR), first quartile, median, third quartile, and maximum (Q3 + 1.5IQR) without outliers. p-values are calculated using the two-tailed Mann-Whitney-Wilcoxon test. (ns P > 0.05; *P < 0.05; **P < 0.01; ***P < 0.001; ****P < 0.0001). **E.** Vplot showing fragment length on y-axis at the Overlapping, Non-overlapping and AR specific sites, with distance from peak summit on x-axis at 1h and 24 h post signaling. The color of pixel denotes depth and number of loci normalized fragment density. **F.** Profile plot of normalized fragment density of different sized fragments (Large (>300bp), NFR(<120bp) and (Sub and Mono Nucleosome(120-300bp) on Overlapping, Non-Overlapping and AR-Specific sites, with distance from peak summit on x-axis at 1 and 24 h. **G.** IgV snapshot of ERα, ATAC-Seq, H3K27ac, p300, Med1, Pol II, GRO-seq at 1hr and AR at 24 h of E2 signaling on the XBP1 gene locus. **H.** Kernel density plots comparing Pol II, ERα, Med1, p300, H3K27ac and GRO-Seq at 1 h of signaling for ER-AR overlapping peaks and non-overlapping peaks. **I.** GRO-seq time course of the nearest gene at different times post-signaling. The y-axis denotes normalised counts, and the x-axis denotes different times post-signaling. The dark green curve shows genes closest to overlapping peaks, the light green curve shows genes closest to non-overlapping peaks, and blue shows genes closest to AR-specific genes. The curve shows the mean ± 95% confidence interval.

V-plots of ATAC-seq signal at these two categories revealed that, although chromatin accessibility decreased globally at ERα-bound regions from 1 to 24 h, AR-ERα overlapping regions retained significantly greater accessibility than ERα regions that did not acquire AR binding (Fig. 2E). In contrast, AR-only regions showed no appreciable change in accessibility between 1 and 24 h. Consistent with the V-plots, nucleosome profiling further demonstrated that AR-ERα overlapping regions retained greater accessibility at 24 h, whereas accessibility was reduced at non-overlapping ERα regions (Fig. 2F). Overall, the overlapping regions displayed higher ERα, Med1, H3K27ac, p300, Pol II, ATAC-seq, and GRO-seq signals than the non-overlapping sites, consistent with their identity as highly active enhancers (Fig. 2G-H; Supp. Fig. 2A-B). Finally, to evaluate the functional impact of these enhancers on rate of target gene expression, we analyzed the transcriptional output of genes proximal to these regions. Genes associated with AR-ERα overlapping regions exhibited both faster and stronger induction upon treatment compared to non-overlapping regions, whereas those near AR sites alone failed to show significant estrogen-induced activation (Fig. 2I).

We previously reported that a subset of ERα regulatory elements, termed “persistent enhancers”, are essential for robust E2-induced gene transcription (Saravanan et al. 2020). Given the high baseline activity and transcriptional machinery enrichment at AR-ERα shared sites, we asked whether these regions correspond to persistent enhancers. Intersecting these datasets revealed that over 75% of the overlapping peaks matched previously defined persistent sites (Supp Fig. 2C-D). Together, these results suggest that late-phase AR recruitment to ERα enhancers serves as a mechanism to keep highly active, persistent regulatory elements marked and accessible.

Additionally, we investigated whether AR targeting of highly active ERα-bound regions is conserved in other ERα positive cellular system. Analyzing public datasets in T47D cells (Hickey et al. 2021), we similarly observed AR binding at ERα regulatory sites. Consistent with our findings in MCF-7 cells, these overlapping regions in T47D cells displayed stronger ERα and AR occupancy, as well as significantly higher H3K27ac levels compared to non-overlapping ERα sites (Supp Fig. 2E-F). Together, these data indicate that AR consistently occupies high-activity ERα enhancers under estrogen stimulation. We call these active enhancers that at late phases of signaling, lose ERα but gain AR and maintain chromatin accessibility as *“reset”* enhancers.

### Reset enhancers lose FOXA1 at the late phase of signaling but remain accessible

As the ATAC-seq profiling indicated that AR-ERα overlapping enhancers maintain chromatin accessibility at 24 h, coinciding with increased AR recruitment, we asked whether AR directly maintains this late-phase accessibility. To test this, we knocked down AR using siRNA (siAR) and assessed chromatin accessibility via ATAC-seq at 24 h. Surprisingly, AR depletion resulted in increased accessibility across all ERα regions with overlapping enhancers showed a higher gain (Fig. 3A–C; Supp Fig. 3A). As these overlapping enhancers are active persistent enhancers, the TSS and +1 nucleosome of the closest genes exhibited enhanced accessibility. These data suggest that AR maintains an optimal, intermediate level of chromatin accessibility at enhancers during the late phase of signaling. In the absence of AR, this controlled reduction in accessibility is disrupted, resulting in increased chromatin opening at these sites.

**Fig. 3:**
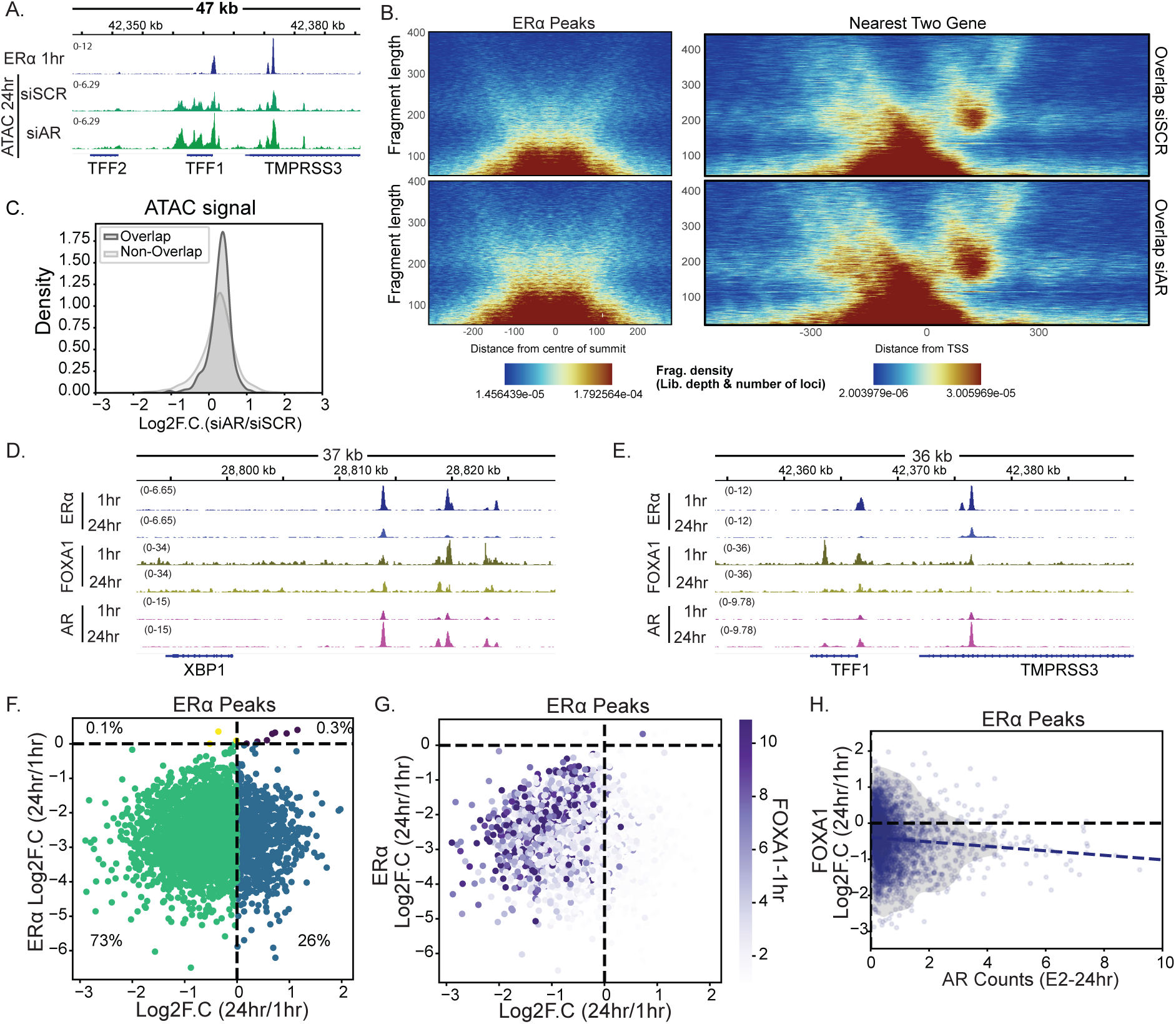
Reset enhancers lose FOXA1 at the late phase of signaling but remain accessible. **A.** IgV snapshot of ERα binding at 1h and ATAC-Seq upon siAR at 24h E2 signaling on the TFF1 gene locus. **B.** Vplot showing fragment length on y-axis at the Overlapping and TSS of nearest two genes with distance from peak summit and TSS respectively on x-axis at 1h and 24 h post signaling in siSCR and siAR. The color of pixel denotes depth and number of loci normalized fragment density. **C.** Kernel Density plot of overlapping and non-overlapping peaks. The x-axis denotes log2FC.(siAR/siSCR) of ATAC-seq signal at 24 h and the y-axis denote kernel density. **D.** IgV snapshot of ERα, AR, FOXA1 at 1/24h E2 signaling on the XBP1 gene locus. **E.** IgV snapshot of ERα, AR, FOXA1 at 1/24h E2 signaling on the TFF1 gene locus. **F.** Scatterplot depicting ERα and FOXA1 signal on ERα peaks at 1 and 24 h of E2 signaling. The x-axis denotes log2FC.(24h/1h) FOXA1 signal and the y-axis denotes log2FC.(24h/1h) ERα signal. Different colors indicate different categories, and the percentages show the total number of peaks in each category. **G.** Scatterplot depicting ERα and FOXA1 signal on ERα peaks at 1 and 24 h of E2 signaling. The x-axis denotes log2FC.(24h/1h) FOXA1 signal, and the y-axis denotes log2FC.(24h/1h) ERα signal. The purple hue denotes the FOXA1 signal at 1h. **H.** Scatterplot of FOXA1 and AR signal on ERα peaks. The y-axis denotes log2FC.(24h/1h) FOXA1 signal and the x-axis show the AR signal at 24 h of E2. The grey hue denotes different levels of contour. The purple dashed line indicates the line of best fit.

To identify potential factors that might be responsible for increased accessibility, we performed differential motif enrichment analysis comparing overlapping and non-overlapping ERα regions. This revealed a significant enrichment of Forkhead box (FOX) family motifs within overlapping regions (Supp Fig. 3B–C). As FOXA1 is a well-established lineage-defining pioneer factor required for ERα binding on regulatory elements (Carroll et al. 2005, 2006, Lupien et al. 2008, Hurtado et al. 2011, Zaret and Carroll 2011), we focused on its involvement.

To determine whether persistent accessibility after AR knockdown correlates with FOXA1 occupancy, we stratified all ERα sites into five quintiles based on post-knockdown accessibility, from Bin 1 (lowest) to Bin 5 (highest) (Supp Fig. 3D–G). Notably, the most accessible group (Bin 5) exhibited the highest gain in accessibility upon ARKD, which also showed strongest FOXA1 binding during peak signaling (1h). Additionally, it was also strongly enriched for AR-ERα overlapping regions (Supp Fig. 3F–G). Together, these findings indicate that the subset of ERα enhancers with the highest initial FOXA1 occupancy during peak signaling (1h) are the same enhancers that show reduced accessibility by 24h, while still retaining higher accessibility than non-overlapping enhancers.

In prostate cancers, FOXA1 and AR engage in a complex, context-dependent relationship wherein FOXA1 facilitates AR chromatin recruitment, with FOXA1 (FKHD) motifs frequently co-occurring adjacent to ARE motifs Supp Fig. 3H (Gao et al. 2003, Sahu et al. 2011, Jin et al. 2014, Robinson et al. 2014, Pomerantz et al. 2015, Vorontsov et al. 2024). In addition, AR binding sites in breast cancer cells is shown to be abundant for FOXA1 motifs (Robinson et al. 2011). This close spatial arrangement prompted us to ask whether FOXA1 binding itself is dynamically altered at overlapping regions during estrogen signaling.

To test this, we performed FOXA1 CUT&RUN at 1h and 24h post-E2 exposure. FOXA1 occupancy declined both globally and specifically at ERα-bound regions, with AR-ERα overlapping sites showing a significantly greater loss than non-overlapping regions (Fig. 3D-E; Supp Fig. 3I-J, M). Comparing occupancy fold changes between 1 h and 24 h revealed that nearly 73% of ERα-bound sites experienced a concurrent decrease in both ERα and FOXA1 (Fig. 3F). The magnitude of this FOXA1 loss scaled directly with its initial 1h occupancy level (Fig. 3G).

Crucially, when we tracked whether sites losing FOXA1 corresponded to late-phase AR recruitment, we found that regions exhibiting FOXA1 loss, gained the highest levels of AR occupancy at 24 h (Fig. 3H; Supp Fig. 3K-L). This inverse dynamic strongly supports a model in which AR acts as a placeholder at these high FOXA1-active enhancers during the late phase of signaling.

These findings were striking in two respects. First, FOXA1 showed decreased binding at the late phase of signaling despite being a lineage-defining pioneer factor (Lupien et al. 2008). This pattern is not typically expected of lineage-defining transcription factors, which are generally considered stable rather than dynamic occupants of enhancers and other genomic regions (Glont et al. 2019).). Second, AR, despite lacking pioneer activity, occupies FOXA1-bound regions and helps maintain their chromatin accessibility as a placeholder. Yet AR loss increases FOXA1 occupancy and reopens these enhancers, indicating that AR actively competes with FOXA1 to constrain, rather than fully close, chromatin at these sites, consistent with an active, competitive handoff between the two factors, rather than a passive, sequential one. Notably, our culture conditions did not include exogenous testosterone (the canonical AR ligand); intracellularly, androgens are converted to estrogens, not the reverse (Payne and Hales 2004).

### AR resets ERα enhancers by evicting FOXA1

We next asked if the loss of FOXA1 is facilitated by direct AR binding at the late phase of signaling. We profiled FOXA1 binding upon AR knockdown and observed that FOXA1 occupancy increased at the late phase of signaling, with a greater increase at overlapping regions (Fig. 4A-B, and Supp Fig. 4A, D, E). Additionally, increased occupancy correlated well with chromatin accessibility at these sites after AR knockdown (Fig 4C and Supp Fig. 4B-C). Total FOXA1 protein levels remained unchanged following AR knockdown after 24h of E2, indicating that AR depletion does not affect FOXA1 protein abundance (Supp Fig. 4F). The data suggest that AR potentially evicts FOXA1 to reset ERα-enhancers at the late phase of signaling.

**Fig. 4:**
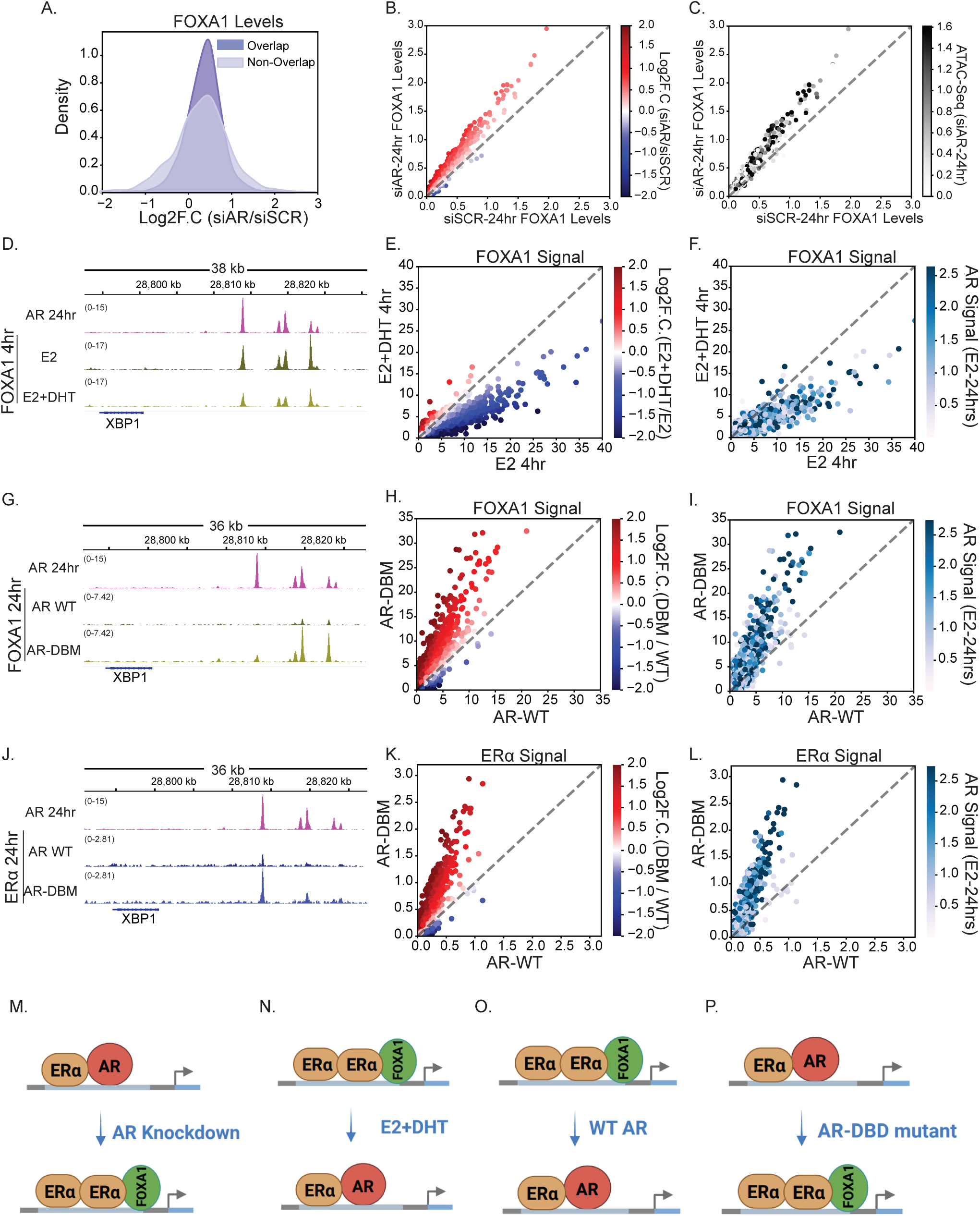
AR resets ERα enhancers by evicting FOXA1. **A.** Kernel Density plot of overlapping and non-overlapping peaks. The x-axis denotes log2FC. (siAR/siSCR) FOXA1 signal at 24 h of E2, and the y-axis denotes kernel density. **B.** Scatterplot of FOXA1 signal on ERα peaks in siSCR and siAR. The y-axis denotes the FOXA1 signal in siAR, and the x-axis denotes the signal in siSCR. The colour of the spot denotes log2FC. (siAR/siSCR) FOXA1 signal at 24 h of E2. **C.** A scatterplot is plotted in the same manner as B. The colour indicates the ATAC-seq signal in siAR at 24 h of E2. **D.** IgV snapshot of AR at 24 h of E2 signaling and FOXA1 signal at 4 h of E2 and E2+DHT treatment on the XBP1 gene locus. **E.** Scatterplot of FOXA1 signal on ERα peaks in E2 and E2+DHT. The y-axis denotes the FOXA1 signal in E2+DHT, and the x-axis denotes the signal in E2. The colour of the spot denotes log2FC. (E2+DHT/E2) FOXA1 signal at 4 h of E2. **F.** Scatterplot is plotted in the same manner as E. The colour indicates the AR signal at 24 h of E2. **G.** IgV snapshot of AR at 24 h of E2 signaling and FOXA1 signal at 24 h upon WT-AR and AR-DBM (DNA-binding mutant) overexpression on the XBP1 gene locus. **H.** Scatterplot of FOXA1 signal on ERα peaks in WT-AR and DBM-AR overexpression. The y-axis denotes the FOXA1 signal in DBM-AR, and the x-axis denotes the signal in WT-AR. The colour of the spot denotes log2FC. (DBM/WT) FOXA1 signal at 24 h of E2. **I.** A scatterplot is plotted in the same manner as H. The colour indicates the AR signal at 24 h of E2. **J.** IgV snapshot of AR at 24 h of E2 signaling and ERα signal at 24 h upon WT-AR and DBM-AR (DNA-binding mutant) overexpression on the XBP1 gene locus. **K.** Scatterplot of ERα signal on ERα peaks in WT-AR and DBM-AR overexpression. The y-axis denotes the ERα signal in AR-DBM, and the x-axis denotes the signal in WT-AR. The colour of the spot denotes log2FC. (DBM/WT) ERα signal at 24 h of E2. **L.** Scatterplot is plotted in the same manner as K. The colour indicates the AR signal at 24 h of E2. **M.** A schematic showing the effect of siAR on the FOXA1 and ERα binding. **N.** A schematic showing the effect of combined E2 plus DHT treatment on the FOXA1 binding. **O.** A schematic showing the effect of overexpression of WT-AR on the FOXA1 and ERα binding. **P.** A schematic showing the effect of overexpression of the AR DNA-binding mutant on FOXA1 and ERα binding.

The intriguing, complex relationship between FOXA1 and liganded-AR in prostate cancer is known (Jin et al. 2014), but unclear. However, this relationship, to our knowledge, is unknown in breast cells, and that too under estrogen stimulation. Recently, AR was shown to sequester coactivators away from ERα-target genes toward AR-regulated genes upon stimulation with DHT (an AR agonist) (Hickey et al. 2021). Although our culture conditions lacked additional DHT, we observed AR binding at ERα-targeted enhancers and loss of FOXA1 during the late phase of signaling. We therefore asked whether adding along DHT to estrogen-stimulated cells, thereby increasing liganded AR at these enhancers, would further enhance FOXA1 loss relative to estrogen alone. Towards this, we profiled FOXA1 upon estrogen or estrogen along with DHT stimulation for 4 h and found FOXA1 loss at the majority of ERα-bound regions, with overlapping regions exhibiting greater loss than non-overlapping regions in estrogen along with DHT as compared to estrogen alone (Fig. 4 D-E and Supp Fig. 5A, E). The loss of FOXA1 was positively correlated with AR occupancy at these regions at the late phase of signaling (Fig. 4F and Supp Fig. 5A, H). The data suggest that AR displaces FOXA1 under estrogen or testosterone stimulation.

**Fig. 5:**
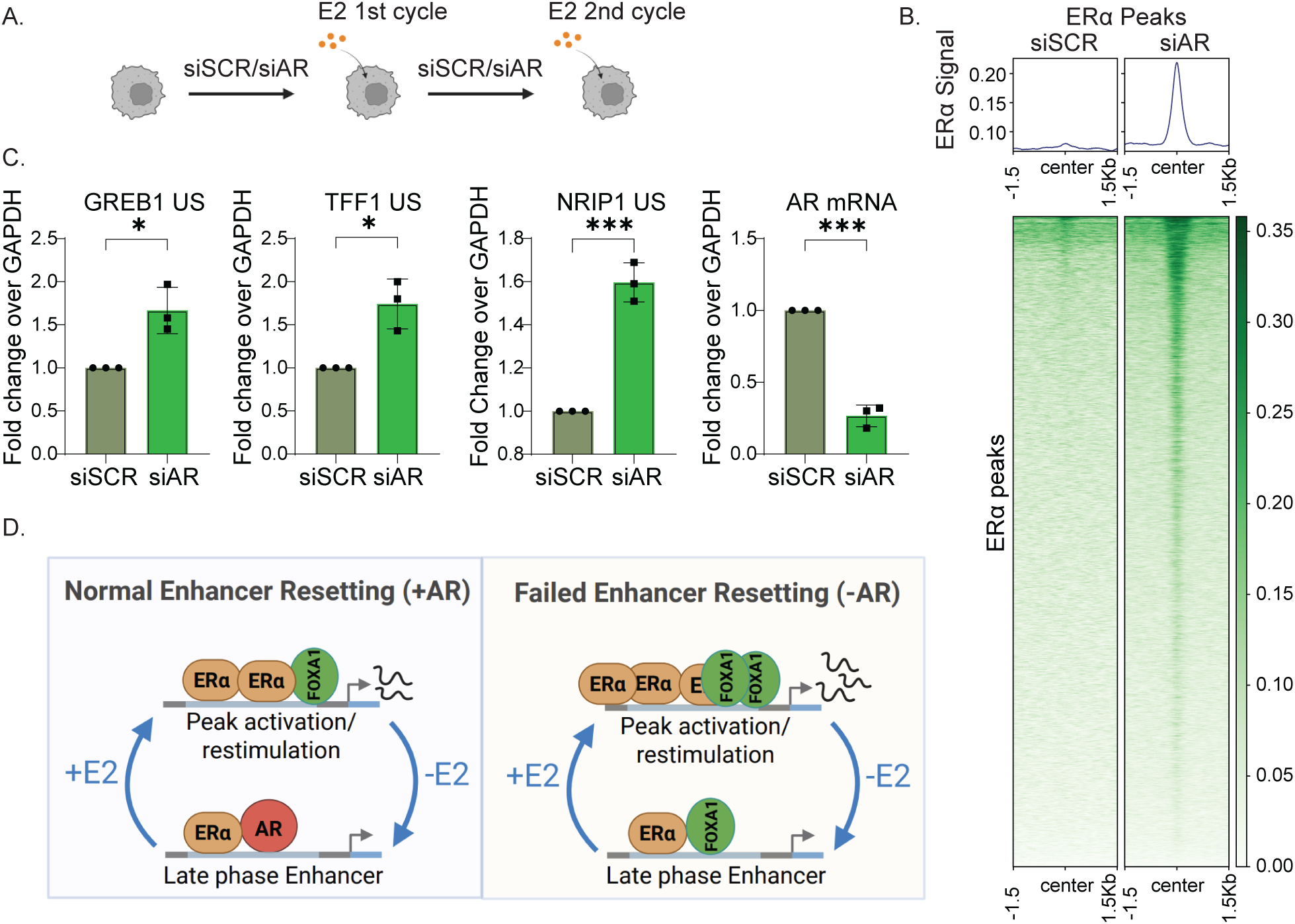
Absence of AR-mediated enhancer-resetting hyperactivates E2-regulated genes during the next cycle of estrogen stimulation. **A.** Schematic of re-signaling treatment upon siAR. **B.** Heatmap showing ERα signal on ERα peaks at 1h of E2 treatment upon re-signaling in siSCR and siAR. **C.** qPCR of E2-responsive genes (TFF1, GREB1 and NRIP1) and AR as a control 1h of E2 in siSCR and siAR in re-signaling. **D.** The schematic shows a model of the study, which shows that during the late phase of estrogen signaling, AR resets ERα enhancers to prevent hyperactivation of the estrogen-mediated transcriptional program. The absence of AR fails to reset the enhancers, which results in higher FOXA1 and ERα DNA binding and leads to a higher estrogen-mediated transcriptional response upon the next cycle of estrogen stimulation.

We then asked whether direct DNA binding by AR is required for FOXA1 displacement. To test this, we generated a DNA-binding mutant of AR (DBM-AR) by introducing an alanine-to-aspartate substitution at amino acid position 573 (A573D) (Farla et al. 2005, Antony et al. 2014). We validated that DBM-AR failed to bind the GREB1 enhancer, a representative late-resetting target (Supp Fig. 5B). We then performed FOXA1 ChIP-seq following overexpression of wild-type AR (WT-AR) or DBM-AR under estrogen stimulation. WT-AR expression resulted in low FOXA1 occupancy on overlapping regions suggesting, high AR levels at late phase of signaling might be the reason of FOXA1 eviction (Fig 4G, H). Further, DBM-AR caused an increase in FOXA1 occupancy across ERα-bound regions at 24 h (Fig. 4G-H; Supp Fig. 5C), with the higher gains occurring at AR-ERα overlapping regions (Supp Fig. 5F). Moreover, this gain in FOXA1 binding correlated positively with late-phase AR occupancy (Fig. 4I; Supp Fig. 5I). Collectively, these results demonstrate that high levels of AR and its direct DNA binding are required at late phase to displace FOXA1 and reset enhancers.

Since FOXA1 is a pioneering factor and its modulation on chromatin is linked to ERα binding in the genome (Hurtado et al. 2011), we asked whether FOXA1 gain upon DBM-AR overexpression w.r.t to WT-AR, in turn affects ERα occupancy. ERα mirrored the FOXA1 gain upon DBM-AR overexpression as revealed by ERα ChIP-seq (Fig. 4J-K and Supp Fig. 5D, G). The increase in ERα occupancy was also positively correlated with the AR binding at these regions at the late phase of signaling (Fig. 4L and Supp Fig. 5J). Total ERα protein levels remained unchanged following overexpression of WT-AR and DBM-AR after 24 h of E2, indicating that overexpression of AR does not affect ERα protein levels (Supp Fig. 5K). Overall, the data suggest that altering AR occupancy, by increasing it with Estrogen, DHT or disrupting it by AR-DBM, produces opposite effects on FOXA1 occupancy and that these changes extend from FOXA1 to ERα (Fig. 4M-P).

### Absence of AR-mediated enhancer resetting hyperactivates E2-regulated genes during the next cycle of estrogen stimulation

We hypothesized that if AR fails to reset enhancers, FOXA1 will be present at the late phase of signaling; consequently, upon the next round of estrogen stimulation, these enhancers will gain more ERα, contributing to higher gene expression. To test this, we knocked down AR, treated cells with E2 for 24h to perturb resetting, and then stripped the cells of hormone for 72h while maintaining low AR levels through a second round of AR knockdown. Cells were then restimulated with estrogen for 1 h (next round of stimulation) (Fig. 5A).

We first assessed ERα binding by ChIP-seq in the next round of E2 signaling and found its occupancy increased relative to the scr siRNA control (Fig. 5B). We next asked whether the enhanced ERα binding was reflected in higher expression of E2-regulated genes. We tested the expression of representative classical E2-regulated genes (TFF1, NRIP1, and GREB1) whose enhancers exhibited resetting at the late phase of signaling. The nascent transcripts of these genes exhibited higher expression as compared to the scr siRNA control (Fig. 5C). Together, these data suggest that AR resets enhancers by evicting FOXA1 during the late phase of signaling to prevent hyperactivation of the estrogen-responsive transcriptional program upon subsequent ligand exposure (Fig. 5D).

## Discussion

Estrogen-dependent transcription is inherently cyclical where acute activation is followed by chronic phase. While extensive work has established how extracellular cues activate enhancers to initiate transcription, considerably less is known about how active enhancers are restored to a competent state for future rounds of stimulation. Here, we identify enhancer resetting as a mechanistically distinct phase of the transcription cycle. Using estrogen signaling as a model, we show that highly active ERα enhancers undergo an ordered transition in which FOXA1 is actively displaced from chromatin by AR, enhancer activity is terminated to close the enhancers but AR maintains limited accessibility (Fig. 2 and 3). Failure of this resetting process results in excessive enhancer activity and hyperactivation of estrogen-responsive genes during subsequent hormone exposure (Fig. 5). These findings establish that enhancer activation is coupled to an equally important decommissioning program that preserves transcriptional fidelity across repeated signaling cycles.

Estrogen signaling has long been understood as a pulsatile system: ERα binds chromatin, mostly enhancers, recruits coactivators, drives transcription, and is then cleared by proteasomal degradation, resetting the system for the next stimulus. What has been missing from this picture is a mechanistic account of how the enhancers themselves, not just the receptor, are restored to a poised, unbiased state between signaling cycles. Our data identify AR as the missing link in this cycle: a receptor best known for opposing ERα occupancy is repurposed, as an enhancer-resetting factor that acts by evicting the pioneer factor FOXA1.

### A non-canonical role of AR under estrogen stimulation

The prevailing model of AR action in ERα+ breast cancer is built almost entirely on liganded AR, where DHT-bound AR competes with ERα for genomic binding sites, redistributing coactivators such as p300 and SRC-3 away from ERα target genes, and redirecting ERα toward androgen response elements (Hickey et al. 2021). This framework, while well supported, does not explain how AR could exert tumor-suppressive, enhancer-protective effects in a tissue context where androgen availability is low and where intracellular steroidogenesis favors conversion of androgens to estrogens rather than the reverse. Our data reveal a second, ligand-independent mode of AR action: as ERα is degraded over the course of a E2 signaling, AR itself accumulates in the nucleus and progressively occupies a defined subset of ERα-bound enhancers (Fig. 1). This occupancy is transcriptionally silent as transcription of genes remains at pre-stimulation expression levels at late phase of signaling, yet it is functionally essential, since its removal by siRNA licenses FOXA1’s continuous occupancy, contributing to heightened ERα occupancy and re-activation of the same loci upon subsequent stimulation (Fig. 4, 5). In other words, AR’s tumour-suppressive activity may not be reducible to a single mechanism but may instead comprise (at least) two temporally and mechanistically distinct programs: acute antagonism when liganded, and chronic, silent gatekeeping under extremely low to absent androgens as ligand. This distinction reframes AR less as a competitor for the ERα program and more as a placeholder of the enhancer ERα leaves behind.

### FOXA1 dynamics challenge the static pioneer-factor paradigm

FOXA1 is conventionally treated as a stable, lineage-defining pioneering factor that licenses ERα binding but is not itself subject to large, cyclical changes in occupancy. Our observation that FOXA1 occupancy decreases over the E2 signaling cycle, that it is displaced from chromatin as AR rises and that it persists when AR is depleted, argues that pioneer factor occupancy at a subset of enhancers is more dynamic and more contested than currently appreciated (Fig. 3, 4). Critically, the sites that lose the most FOXA1 are precisely the sites that gain the most AR, and reversing AR occupancy (by knockdown or by a DNA-binding-dead AR mutant) restores FOXA1, while increasing AR occupancy (via DHT) deepens the loss (Fig. 4).

### Decoupling “open” from “occupied”: a handoff model of enhancer accessibility

A central puzzle our data raise, and partially resolve, is how chromatin accessibility is decoupled from the identity of the transcription factor sitting on the motif. FOXA1 departs these enhancers by 24h, accessibility is reduced, but loss of AR increases the accessibility (Fig. 1I). This is best explained by a handoff model: FOXA1 performs the pioneering, nucleosome-displacing work required to open the enhancer in the first place, but once open, consistent accessibility is maintained by continuous occupancy of the motif first by FOXA1, then by AR. This distinguishes two separable functions long conflated under “pioneer activity”: the ability to invade nucleosomal DNA (a property specific to true pioneer factors like FOXA1) versus the ability to maintain an already-accessible motif against nucleosome reformation (a property any DNA-binding factor occupying that site, including non-pioneer AR, can provide). Under this model, AR’s occupancy is not incidental to accessibility; it is a stopgap: a way of keeping the site FOXA1-free and transcriptionally quiet without allowing the enhancer to fully collapse and lose its “memory” of prior activity.

### Enhancer resetting as a rheostat against hyperactivation

The functional consequence of this handoff is perhaps the most clinically resonant finding: enhancers that fail to undergo AR-mediated FOXA1 eviction do not simply retain a passive memory of prior signaling; they hyperactivate upon restimulation, producing both elevated ERα rebinding and elevated nascent transcription at canonical E2 target genes (Fig. 5). This suggests that the estrogen response is not simply “on” or “off” between pulses, but is actively normalized by AR-dependent resetting, and that loss of this resetting function constitutes a distinct route to pathological hyperactivation of ER signaling, conceptually related to, but mechanistically separate from, loss of ERα degradation or coactivator overexpression, the more commonly studied routes to hormone hypersensitivity in breast cancer. Given that AR expression and activity are frequently altered in the natural history of ERα+ breast cancer, this raises the possibility that a decline in AR’s resetting capacity, rather than only its coactivator-competition capacity, contributes to the transition toward hormone hypersensitivity or endocrine therapy resistance. However, it is yet to be tested.

### Relationship to persistent enhancers and enhancer memory

Our finding that the majority of AR-occupied, FOXA1-competitive enhancers correspond to previously defined persistent enhancers, the enhancers that remain open while other ERα enhancers lose accessibility at the end of signaling (Saravanan et al. 2020) (Supp Fig. 2C,D), suggests these two phenomena are mechanistically linked facets of the same underlying biology: the same enhancer subset that is primed to respond fastest and most robustly to hormone is also the subset requiring the most active intervention to reset. This raises a broader principle for enhancer biology: the enhancers with the greatest regulatory potency may be precisely those at greatest risk of runaway reactivation, and may therefore require dedicated, factor-specific resetting machinery rather than passive chromatin closure. AR, in this tissue and hormonal context, appears to be that machinery.

### Limitations and open questions

Our evidence for direct AR-FOXA1 motif competition, while supported by correlative genomic and genetic-perturbation data, would benefit from base-pair-resolution co-occupancy analysis or single-molecule foot-printing to confirm mutual exclusivity at the same allele. Second, the experiments were largely established in MCF-7 and T47-D ERα-positive cell lines; future studies in tissues and ultimately in patient-derived tumors will be necessary to determine the generality of the observation.

## Methods

### Cell culture

MCF-7 cells were obtained from ATCC and cultured in high-glucose DMEM (10569044 Invitrogen) supplemented with 10% fetal bovine serum (Invitrogen 16000044) and 1% penicillin-streptomycin (Gibco 15140122) at 37°C in a humidified incubator containing 5% CO₂. For hormone deprivation experiments, MCF-7 cells were initially seeded in complete DMEM, and the next day, cells were washed with 1× DPBS (14190144 Gibco) and cultured in phenol red-free DMEM (21063045 Invitrogen) supplemented with 5% charcoal-stripped FBS (12676029 Invitrogen). After 72 h of hormone deprivation, cells were treated with 100 nM 17β-estradiol (Sigma-Aldrich, E2758) or vehicle control (ethanol; CAS number 64-17-5, Millipore) for the indicated time points.

For overexpression studies, transfections were performed using Lipofectamine 2000 (Invitrogen, 11668019) according to the manufacturer’s instructions. Transfections were carried out in phenol red-free Opti-MEM reduced serum medium (Invitrogen, 11058021). Six hours post-transfection, the medium was replaced with hormone-depleted stripping medium, and cells were subsequently stimulated with 100 nM 17β-estradiol on day 3.

### Generation of AR DNA-binding mutant plasmids

Wild-type AR (WT-AR) and the A573D DNA-binding mutant (DBM-AR) were generated by PCR amplification using the pLENTI6.3/AR-GC-E2325 plasmid (Addgene #85128) as the template. The primers used for amplification are listed in Appendix Table S1. The PCR-amplified WT-AR and A573D mutant fragments were then cloned into the 3×FLAG-CMV10 vector using NotI (R3189L) and XbaI (R0145S) restriction sites.

### siRNA knockdown experiments

MCF7 cells were seeded, and the next day, culture medium was replaced with stripping medium. AR knockdown was carried out over two consecutive days using AR-specific siRNA (Catalog ID: L-003400-00-0010), with scrambled siRNA (Catalog ID: D-001810-10-05) serving as a negative control. Following knockdown, cells were treated with E2 for the indicated time points.

### Western Blotting

MCF-7 cells were seeded in 6-well plates and, the following day, subjected to hormone deprivation for 72 h. After hormone deprivation, cells were treated with 100 nM 17β-estradiol (E2) or vehicle control for the indicated time points. Following treatment, culture medium was aspirated, and cells were washed three times with ice-cold 1× PBS and harvested by scraping; cell pellets were lysed in RIPA (NaCl 150 mM, Nonidet P-40 1%, Sodium deoxycholate 0.5 %, SDS 0.1%, Tris pH 7.4 50mM) buffer supplemented with 1× protease inhibitor cocktail (Roche 05056489001). Lysates were incubated on ice for 40 minutes, followed by sonication (Bioruptor Diagenode, 10 cycles to ensure complete lysis. Samples were then centrifuged at 12,000 rpm for 12 minutes at 4 °C, and the supernatant was collected. Protein concentration was determined using the Bradford assay (#5000006 Bio-Rad). Then, 6X Laemmli sample buffer was diluted to a final concentration of 1X and added, and samples were denatured by boiling at 100 °C for 10 minutes. Equal amounts of protein were loaded using SDS-PAGE and subsequently transferred onto PVDF (Thermo 88518) membranes. Membranes were blocked with 5% skim milk in TBST at room temperature for 1 h with gentle shaking on an orbital shaker, followed by incubation with an AR (ab108341) and ERα (sc-8002), primary antibodies overnight at 4 °C. After washing with 1× TBST, membranes were incubated with HRP-conjugated secondary antibodies. Blots were washed three times with 1X TBST and developed using the ImageQuant LAS 4000 imaging system.

### Immunostaining

Cells were washed three times with ice-cold 1× PBS and fixed with 4% paraformaldehyde (P6148-500G) for 10 minutes at room temperature (RT). Following fixation, cells were washed twice with 1× PBS and permeabilized with 1× PBS containing 0.3% Triton X-100 (T8787-250ml) for 15 minutes at RT. Cells were then washed twice with 1× PBS. Blocking was performed using 1% bovine serum albumin (PG-2330-100G) in 1× PBS for 15 minutes at RT, followed by two washes with 1× PBS. Primary antibodies against ERα (sc-8002, 1:500) and AR (ab108341, 1:500) were diluted in 1% BSA/1×PBS, and cells were incubated with the primary antibody solution overnight at 4°C. After incubation, cells were washed three times with 1×PBST, and then appropriate fluorophore-conjugated secondary antibodies were diluted 1:500 in 1% BSA/1× PBS and applied to the cells for 1 h at RT. Cells were then washed three times with 1×PBST and stained with DAPI for 2 minutes at RT, followed by three additional washes with 1×PBST. Coverslips were air-dried to remove excess liquid and mounted onto microscope slides using 90% glycerol as a mounting medium. Fluorescently labeled cells were imaged using a confocal laser scanning microscope (FV3000 6-laser system; Olympus) at the CIFF Microscopy Core Facility.

### CUT&RUN

Briefly, 200,000 MCF-7 cells were harvested and washed twice with 1ml wash buffer (20mM HEPES, pH 7.5, 150mM NaCl, 0.5mM spermidine and 1X PIC). Then cells were resuspended in 100 µL wash buffer and incubated with 10 µL activated concanavalin A beads (BP531, BangsLabs) for 5 min, for cells to attach to the beads. Cells were then incubated with 100 µL antibody buffer (20mM HEPES; pH 7.5, 150mM NaCl, 0.5mM spermidine, 0.05% digitonin, 2 mM EDTA and 1X PIC) containing 0.6 µg FoxA1 antibody (#53528S, Cell Signaling Technology) overnight at 4°C. The next day, immobilized cells were washed twice with 1ml digitonin wash buffer (20mM HEPES; pH 7.5, 150mM NaCl, 0.5mM spermidine, 0.05% digitonin). Cells were then incubated with 50 µL pAG-MNase mix prepared by diluting 1.5 µL pAG-MNase (#40366, Cell Signaling Technology) in 50 µL digitonin wash buffer, for 1 h at 4°C. After this incubation, cells were further washed twice with 1 mL digitonin wash buffer and resuspended in 150 µL digitonin wash buffer and chilled on ice for 5 min. Finally, 3 µL of 100 mM ice-cold CaCl2 was added to it to start digestion by keeping it at 4°C for 30min. Finally, the reaction was stopped by adding 150 µL 1X stop buffer (170 mM NaCl, 10 mM EDTA, 2 mM EGTA, 0.05% digitonin, and 50 µg/mL RNase A) and incubating at 37°C for 10min. DNA was purified by phenol:chloroform:isoamyl alcohol (Cat# AM9732) extraction. Sequencing libraries were prepared using the NEBNext Ultra II DNA Library Prep Kit for Illumina NEB #E7645S (New England Biolabs). Libraries were sequenced on a NovaSeq 6000 platform to a depth of approximately 12-15 million reads per sample.

### Chromatin immunoprecipitation

ChIP protocol was followed as previously described in (Saravanan et al., 2020). Cells were crosslinked with 1% formaldehyde (Sigma-Aldrich, F8775) for 10 minutes at room temperature with gentle agitation (60 rpm). Crosslinking was quenched by the addition of glycine to a final concentration of 125 mM and incubated for 5 minutes. Cells were washed three times with ice-cold 1×PBS, scraped into 3 mL 1× PBS, and pelleted by centrifugation at 2,500 rpm for 5 minutes at 4 °C, and the crosslinked cell pellets were stored at −80 °C until further use. Cell pellets were resuspended in L2 nuclear lysis buffer (50 mM Tris-HCl pH 7.4, 1% SDS, 10 mM EDTA pH 8.0) supplemented with 1xProtease inhibitor cocktail (PIC) and incubated on ice for 10 minutes. Chromatin was sheared using a Bioruptor sonicator (Diagenode) with 30-second ON/OFF cycles for 10–15 cycles to obtain DNA fragments ranging from 200–600 bp. Lysates were clarified by centrifugation at 12,000 rpm for 12 minutes at 4°C. For each immunoprecipitation (IP), 100 µg of sheared chromatin was diluted 2.5 times with Dilution Buffer (20 mM Tris-HCl pH 7.4, 100 mM NaCl, 2 mM EDTA pH 8.0, 0.5% Triton X-100) supplemented with 1x protease inhibitor cocktail (PIC). Ten percent of the diluted chromatin was kept as an input control. For each IP, 1 µg of the antibody (FOXA1 (ab55178), ERα (sc-8002), Flag (Sigma F7425) was added, and samples were incubated overnight at 4 °C on a rotating platform. Protein G Dynabeads (Invitrogen, 140004D) were blocked in 1% BSA (in 1× PBS) for 1 h at 4 °C and washed with 1× PBS. 15 ul of blocked beads were added per IP and incubated for 4 h at 4 °C with rotation. Beads were sequentially washed at 4 °C with the following buffers: Wash Buffer I (20 mM Tris-HCl pH 7.4, 150 mM NaCl, 0.1% SDS, 2 mM EDTA pH 8.0, 1% Triton X-100), Wash Buffer II (20 mM Tris-HCl pH 7.4, 500 mM NaCl, 2 mM EDTA pH 8.0, 1% Triton X-100), Wash Buffer III (10 mM Tris-HCl pH 7.4, 250 mM LiCl, 1% NP-40, 1% sodium deoxycholate, 1 mM EDTA pH 8.0), and finally 1× TE buffer (10 mM Tris-HCl pH 8.0, 1 mM EDTA pH 8.0). Chromatin was eluted in 200 µL Elution Buffer (100 mM NaHCO₃, 1% SDS) by incubation at 37°C for 30 minutes in a thermomixer at 1,350 rpm. Eluates were transferred to fresh tubes, supplemented with 14 µL of 5 M NaCl, and incubated overnight at 65 °C to reverse cross-linking. DNA was purified using phenol: chloroform: isoamyl alcohol extraction (Ambion, AM9732) followed by ethanol precipitation. The final DNA pellet was air-dried and resuspended in 50 µL of 1× TE buffer. Samples were submitted for library preparation in NGS facility.

Chromatin immunoprecipitation for AR was performed as described above, with some modifications. A total of around 140 µg of chromatin DNA was used for each immunoprecipitation, and 3 µL of anti-AR antibody (Cell Signaling Technology, Cat. No. #5153 D6F11) was added to each reaction. Following immunoprecipitation, the chromatin was washed twice with Wash Buffer 2, while the Wash Buffer 3 step was omitted. DNA was subsequently purified using a PCR purification column according to the manufacturer’s instructions.

### ATAC-seq

A total of 230,000 cells were harvested by centrifugation at 500 × g for 5 minutes at room temperature (RT). The supernatant was discarded, and the cell pellet was retained. Cells were washed with 1× PBS containing 0.01% BSA and centrifuged again at 500 × g for 5 minutes at 4 °C. The supernatant was removed, and the cell pellet was retained for downstream steps. Cells were resuspended in 115 µL ice-cold ATAC lysis buffer (10 mM Tris-HCl pH 7.4, 10 mM NaCl, 3 mM MgCl₂, 0.1% NP-40) and gently mixed by pipetting and inverting 5-6 times. The suspension was incubated on ice for 1 minute and immediately centrifuged at 500 × g for 10 minutes at 4 °C to isolate nuclei. Nuclei were washed with 500 µL wash buffer (10 mM Tris-HCl pH 7.4, 10 mM NaCl, 3 mM MgCl₂, 0.1% Tween-20) and centrifuged at 500 × g for 10 minutes at 4 °C. The supernatant was discarded, and the nuclei pellet was resuspended in 23.5 µL 1× PBS containing 0.01% BSA. Nuclei number was determined using the Trypan blue assay.

For the tagmentation reaction, 50,000 nuclei were resuspended in 50 µL of tagmentation reaction buffer consisting of 25 µL 2× TD buffer, 22.5 µL 1× PBS (0.01% BSA) containing 50,000 nuclei, and 2.5 µL TDE1 transposase (Illumina Tagment DNA Enzyme and Buffer; REF 20034210). The reaction was incubated at 37 °C for 30 minutes with shaking at 800 rpm in a thermoblock. Tagmented DNA was purified using a MinElute column in a final 20 µL elution buffer (Qiagen, 28004). For library preparation, a total of 8 cycles were given for indexing PCR.

### Profile plot and Vplot generation

For each fragment-size class, fragment centers were determined by reducing each fragment to a single-base genomic interval positioned at its midpoint. Genomic regions of interest were represented by their corresponding summit positions, which were similarly reduced to single-base intervals. A symmetric window extending 600 bp upstream and downstream of each region summit was then generated. Fragment centers overlapping these windows were identified using genomic interval overlap analysis, with strand information ignored.

The position of each fragment center relative to the corresponding region summit was calculated as the difference between the genomic coordinate of the fragment center and the summit coordinate. Only fragments located within ±600 bp of the region center were retained. Relative fragment positions were subsequently binned into 10-bp intervals across the ±600-bp window. Fragment counts within each bin were divided by the total number of regions in the respective region set to obtain the number of fragments per region. To account for differences in sequencing library depth, these values were normalized to one million total fragments in the corresponding library (global RPM normalization), using the total number of fragments as the library-size denominator. This normalization was applied consistently across all fragment-size classes.

Fragment profiles were generated separately for the 1 h, 24 h, scramble (SCR), and AR knockdown (KD) conditions and across different genomic region categories. The resulting profiles were combined for visualization. Fragment density was plotted as a function of distance from the region summit, with the region center represented by position 0 bp. Separate panels were generated for each fragment-size class and genomic region category using a faceted plot, with the y-axis scaled independently for each panel to facilitate comparison of fragment-distribution patterns. V-plots were generated for each condition using vplotR.

### ChIP-seq Analysis

Adapters were first filtered out from raw reads using trimgalore. After filtering, reads were aligned to the hg38 human genome using bowtie2. Then, unaligned, unpaired (for paired-end sequencing), discordant and duplicate reads were removed using samtools. Finally, reads with a quality score less than 30 were filtered out. MACS2 was used to call narrow/broad peaks with default parameters. Peaks consistent between the replicates were used for analysis (n=2). Normalized counts on peaks were calculated by taking the average of both replicates.

### CUT&RUN analysis

Adaptors were removed from raw reads using trim galore with default settings. Reads were then mapped to the human genome (hg38) using bowtie2. Then, unaligned, unpaired (for paired-end sequencing), discordant and duplicate reads were removed using samtools. Finally, reads with a quality score less than 30 were filtered out. For spike-in, reads were aligned to *S. cervisiae* genome (SacCer3) using bowtie2 using following parameters: --end-to-end --very-sensitive --no-overlap --no-dovetail --no-mixed --no-discordant --phred33 -I 10 -X 700. The effective spike-in amount was calculated by scaling the differences in human-aligned reads. The spike-in ratio was calculated by taking the ratios of effective spike-in amount. bamCoverage from the DeepTools suite was used to convert bam to bigwig with --scaleFactor for downsampling samples according to spike-in ratios and CPM normalization.

### RNA Isolation and cDNA Synthesis

Total RNA was extracted using TRIzol reagent (Invitrogen, 15596018) according to the manufacturer’s instructions. Briefly, cells were lysed in 1 mL of TRIzol and incubated on a shaker for 10 minutes at room temperature (RT). Chloroform was added at 1/5th the TRIzol volume, the mixture was inverted and mixed thoroughly, and then centrifuged at 12,000 rpm for 12 minutes at 4 °C. The aqueous phase was carefully collected and transferred to a fresh RNase-free tube. RNA was precipitated by adding an equal volume of isopropanol and incubating at RT for 10 minutes. The RNA was pelleted by centrifugation at 12,000 rpm for 10 minutes, and the supernatant was discarded. The pellet was washed twice with 80% ethanol, air-dried for 15 minutes, and resuspended in RNase-free water. To remove residual genomic DNA, RNA was treated with ezDNase (Invitrogen, 117660) according to the manufacturer’s protocol. For cDNA synthesis, 1 µg of DNase-treated RNA was used as input, and reverse transcription was performed using the SuperScript IV First-Strand Synthesis Kit (Invitrogen, 18091050) following the manufacturer’s instructions.

### GRO-seq analysis

Adapters were first filtered out from raw reads using trimgalore. After filtering, reads were aligned to the hg38 human genome using hisat2. Unaligned reads and reads with a quality score less than 30 were filtered out. featureCounts was used to calculate counts on complete transcripts. EdgeR was used to calculate gene RPKM values. To call the closest gene, the bedtools closest function was used with the -d flag to filter out genes more than 200kb away.

## Data and code availability

The next-generation sequencing (NGS) datasets generated in this study, including CUT&RUN, ChIP-seq, and ATAC-seq, are publicly available at the Gene Expression Omnibus (GEO).

## Acknowledgments

We gratefully acknowledge the support of the Department of Atomic Energy, Government of India (Project No.12-R&D-TFR-5.04-0800) and intramural funds from NCBS-TIFR (DN). We also thank the DBT/Wellcome Trust India Alliance for funding support (IA/S/23/1/506749; DN). AHN was supported by CSIR-SRF fellowship. NS acknowledges DBT for support. RM, SI, PK and DS are supported by the NCBS/TIFR graduate program.

## Author contributions

AHN and DN conceived the project, designed the experiments, and wrote the manuscript. AHN performed most experiments with help from RM, SI, DS and PK. RM compiled Figures, and performed most of the NGS data analysis. SN, RNV, SI helped with NGS data analysis. All authors read and edited the manuscript.

---

**Supplementary Figure. 1:**
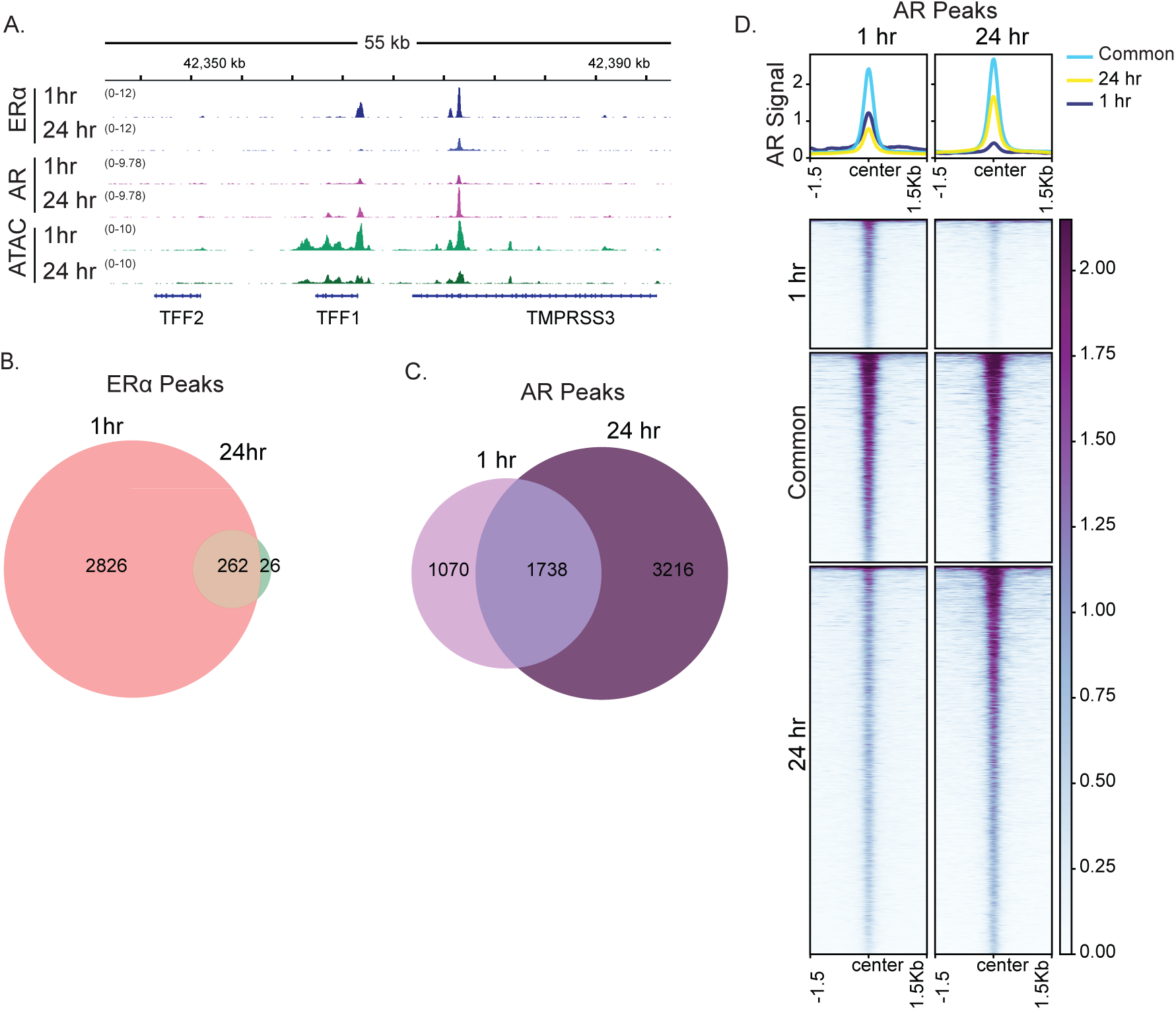
AR accumulates in the nucleus at the late phase of E2 signaling. **A.** IgV snapshot of ERα, AR and ATAC-seq signal at 1 and 24 h of E2 signaling on the TFF1 gene locus. **B.** Venn diagram showing 1hr specific/common/24hr specific ERα peaks. **C.** Venn diagram showing 1hr specific/common/24hr specific AR peaks. **D.** Heatmap showing the AR signal at 1 and 24 h of E2 signaling on 1hr specific/common/24hr specific AR peaks.

**Supplementary Figure. 2:**
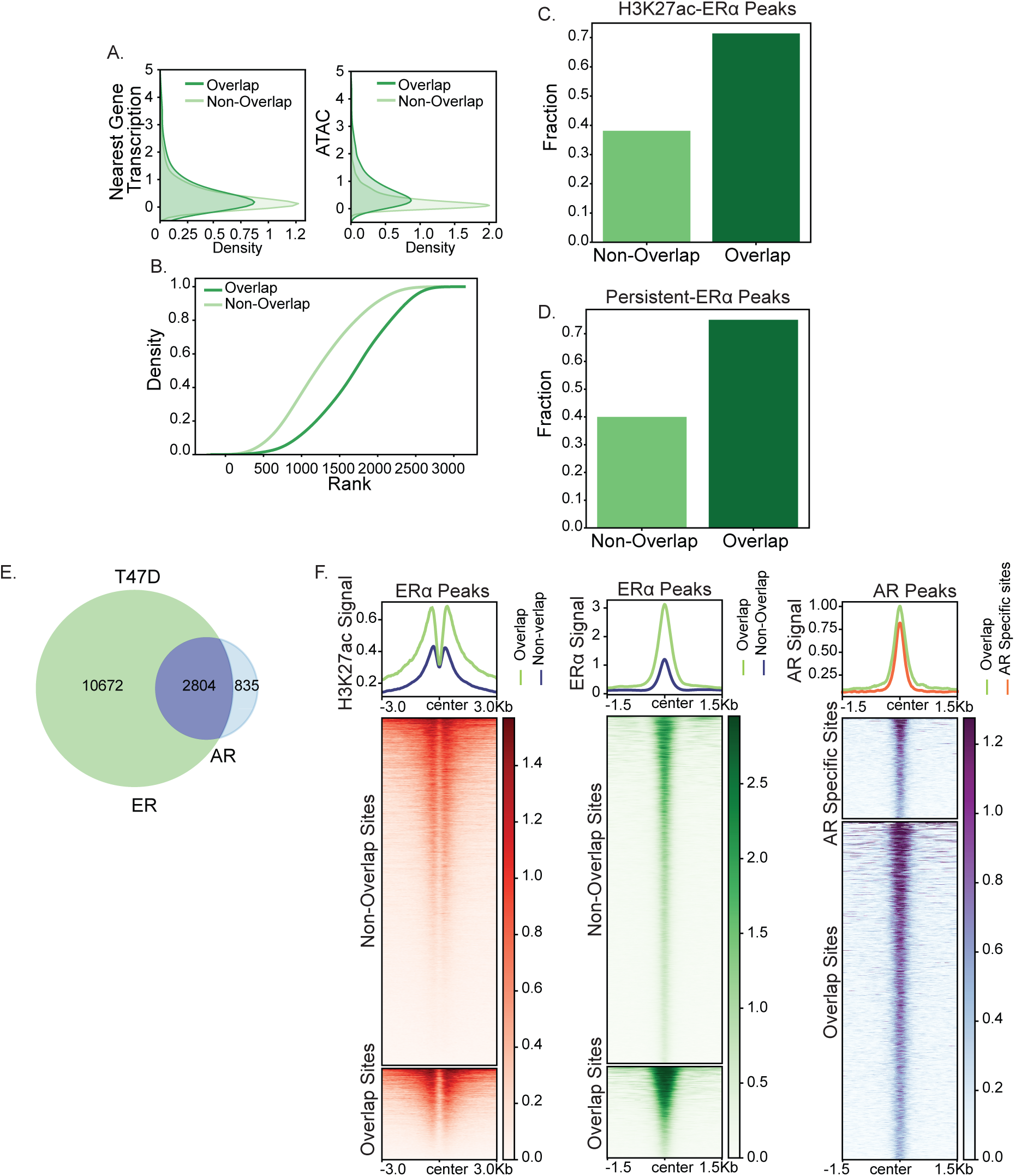
Overlapping ERα sites show active enhancer marks. **A.** Kernel density plots comparing ATAC-Seq and Nearest Gene Transcription at 1 h of signaling for ER-AR overlapping peaks and non-overlapping peaks. **B.** Cumulative density plot of mean rank for Overlapping and Non-Overlapping peaks. **C.** Barplot of the fraction of non-promoter peaks showing H3K27ac enrichment. The y-axis denotes the fraction of peaks. The x-axis denotes the two categories, Overlap and Non-overlap. **D.** Barplot of the fraction of peaks overlapping with persistent enhancers. The y-axis denotes the fraction of peaks. The x-axis denotes the two categories, Overlap and Non-overlap. **E.** Venn diagram showing the intersection of ERα peaks and AR peaks in T47D cells. **F.** Heatmap showing the H3K27ac (left panel), ERα (middle panel), and AR (right panel) counts on AR: ER Overlapping and Non-Overlapping Peaks in T47D cells.

**Supplementary Figure. 3:**
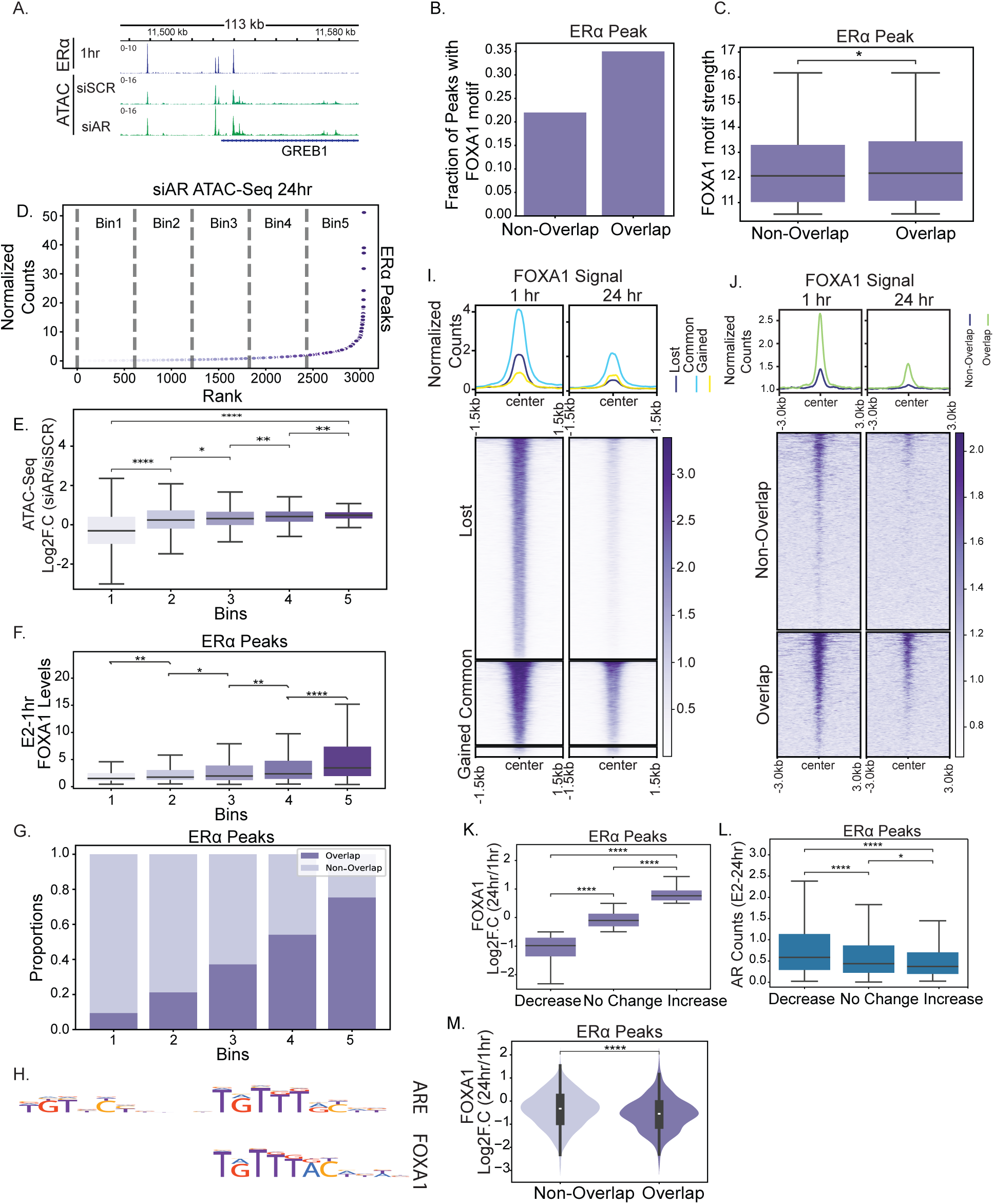
Sites gaining accessibility upon ARKD show enrichment of FOXA1 motif. **A.** IgV snapshot of AR at 24 h of E2 signaling and ATAC-seq signal at 24 h in si-SCR and si-AR on the GREB1 gene locus. **B.** Barplot of the fraction of peaks showing the FOXA1 motif in overlapping and non-overlapping peaks. **C.** Boxplot of FOXA1 motif strength calculated from FIMO in overlapping and non-overlapping peaks. The boxplots depict the minimum (Q1-1.5IQR), first quartile, median, third quartile, and maximum (Q3 + 1.5IQR) without outliers. p-values are calculated using the one-tailed Mann-Whitney-Wilcoxon test with the assumption that non-overlapping is smaller than Overlapping. (ns P > 0.05; *P < 0.05; **P < 0.01; ***P < 0.001; ****P < 0.0001). **D.** Scatterplot of ATAC-seq signal at 24 h in siAR. The x-axis denotes the ranks based on ATAC-seq signal, and the y-axis denotes the ATAC-seq signal in siAR. Dashed lines indicate the different bins, ordered from 5-1 by decreasing ATAC-seq signal. **E.** Boxplot of ATAC-Seq gain upon ARKD in different bins ordered from 5-1 by decreasing ATAC-seq signal. The y-axis denotes the fold change and the x-axis denotes ATAC-seq bins. **F.** Boxplot of FOXA1 signal at 1hr of E2, on ERα peaks binned into 1-5 based on D. **G.** Stacked barplot of the fraction of overlapping and non-overlapping ERα peaks in different bins based on D. **H.** The canonical AR palindromic response element (top) and a composite FOXA1 binding motif (bottom). **I.** Heatmap showing the FOXA1 normalized counts at 1 and 24 h of E2 on FOXA1 peaks lost at 24 h, common between 1 and 24 h, and gained at 24 h. **J.** Heatmap showing the FOXA1 normalized counts at 1 and 24 h of E2 on AR:ER Overlapping and Non-overlapping Peaks. **K.** Boxplot of log2FC. FOXA1 (24hrs/1hr) on ERα peaks shows an increase, no change, and a decrease in FOXA1. The y-axis shows the fold change values, and the x-axis denotes the different conditions. **L.** Boxplot of AR levels at 24 h on ERα peaks showing increase, no change, and decrease based on log2FC. FOXA1 (24hrs/1hr). The y-axis shows the AR levels at 24 h, and the x-axis denotes the different conditions. **M.** Violin plot showing FOXA1 loss quantified by log2FC. FOXA1 (24hrs/1hr) on overlapping and non-overlapping peaks. The y-axis shows fold change, and the x-axis denotes peak categories.

**Supplementary Figure. 4:**
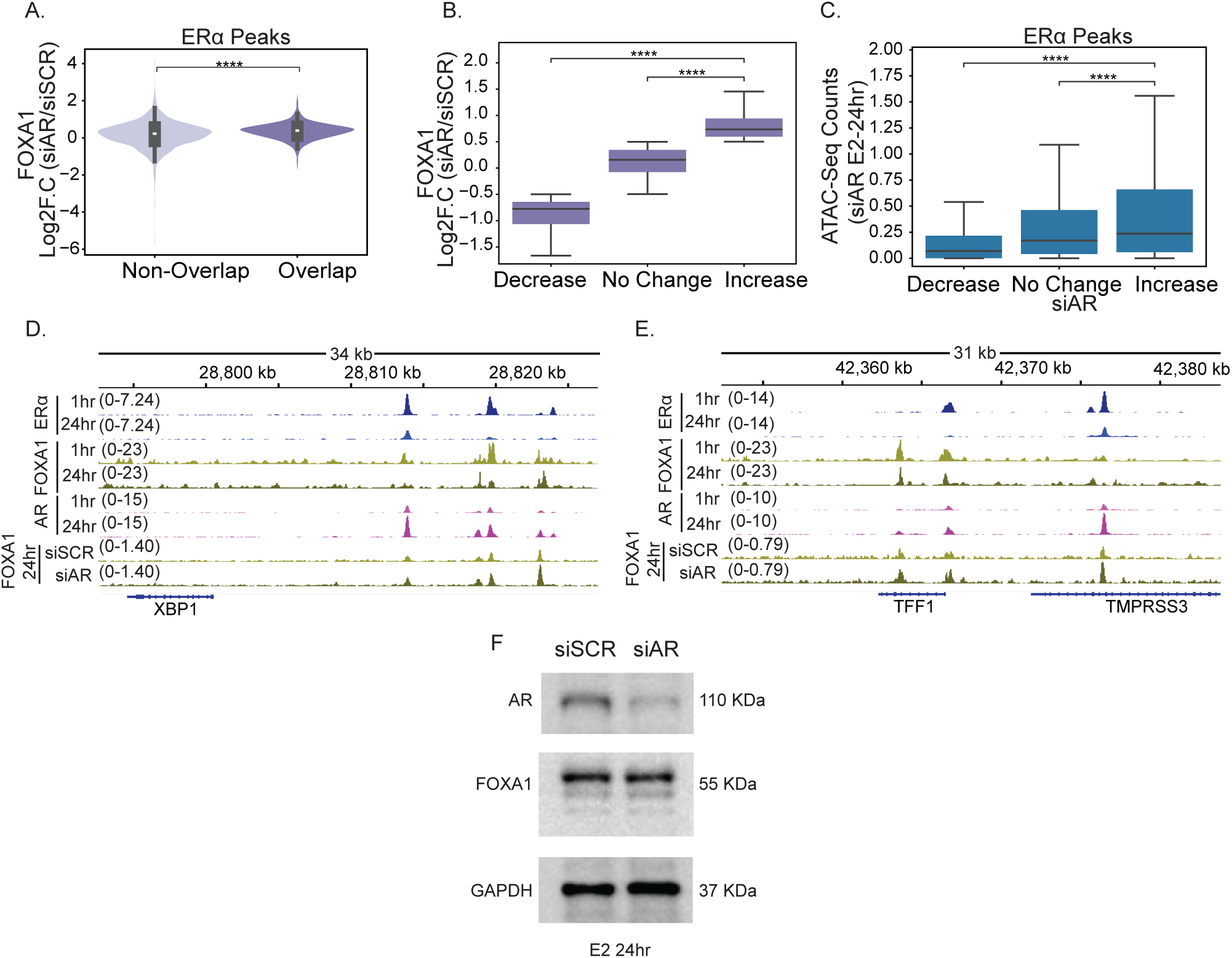
FOXA1 occupancy increases on ERα sites upon ARKD. **A.** Violin plot showing FOXA1 gain quantified by log2FC. FOXA1 (siAR/siSCR) on overlapping and non-overlapping peaks. The y-axis shows fold change, and the x-axis denotes the category of peaks. **B.** Boxplot of log2FC. FOXA1 (siAR/siSCR) on ERα peaks shows an increase, no change, and a decrease upon AR knockdown. The y-axis shows the fold change values and the x-axis denotes the different conditions. **C.** Boxplot of ATAC-seq counts at 24 h in siAR on ERα peaks showing increase, no change, and decrease based on log2FC. FOXA1 (siAR/siSCR). The y-axis shows the AR levels at 24 h, and the x-axis denotes the different conditions. The boxplots depict the minimum (Q1-1.5IQR), first quartile, median, third quartile, and maximum (Q3 + 1.5IQR) without outliers. p-values are calculated using the one-tailed Mann-Whitney-Wilcoxon test with the assumption that Non-Overlapping is smaller than Overlapping. (ns P > 0.05; *P < 0.05; **P < 0.01; ***P < 0.001; ****P < 0.0001). **D.** IgV snapshot of ERα, AR, and FOXA1 at 1/24hr E2 signaling and FOXA1 signal at 24hr in si-SCR and upon si-AR on the XBP1 gene locus. **E.** IgV snapshot of ERα, AR, and FOXA1 at 1/24hr E2 signaling and FOXA1 signal at 24hr in siAR and upon siAR on the TFF1 gene locus. **F.** Western blot of MCF7 cell lysates showing FOXA1 and AR protein levels at 24 h of E2 treatment following AR knockdown using siAR. The blot confirms efficient AR knockdown, while total FOXA1 protein levels remain unchanged upon AR depletion. GAPDH was used as a loading control.

**Supplementary Figure. 5:**
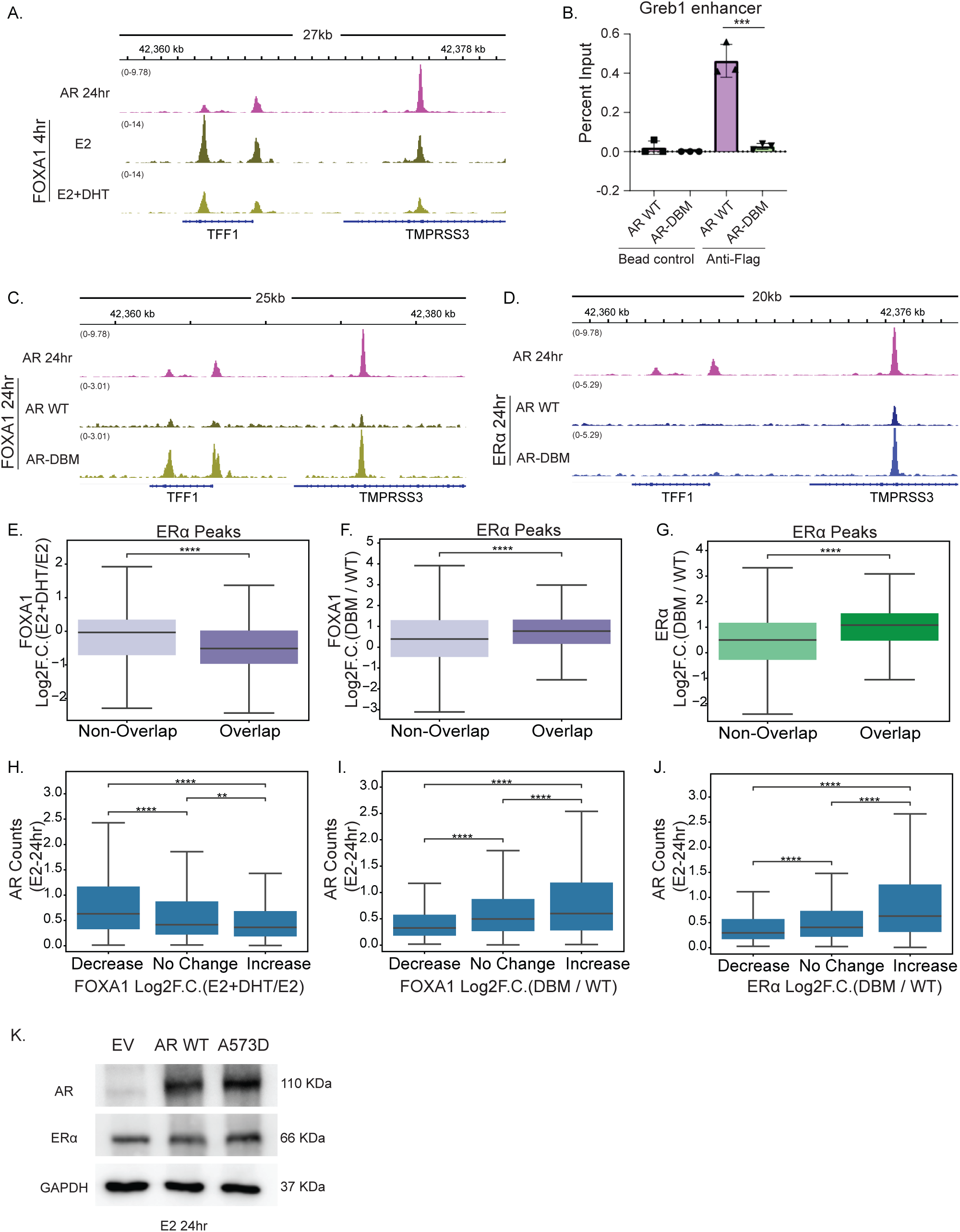
AR-FOXA1 competition shapes ERα program. **A.** IgV snapshot of AR at 24 h of E2 signaling and FOXA1 signal at 4 h of E2 or E2+DHT on the TFF1 gene locus. **B.** ChIP-qPCR of anti-Flag (AR) on the GREB1 enhancer for WT-AR and DBM-AR. The y-axis shows percent input, and the x-axis shows the respective IP and Bead control. (n=3) ns P > 0.05; *P < 0.05; **P < 0.01; ***P < 0.001; ****P < 0.0001. **C.** IgV snapshot of AR at 24 h of E2 signaling and FOXA1 signal at 24 h of E2 in WT-AR or AR-DBM overexpression on the TFF1 gene locus. **D.** IgV snapshot of AR at 24 h of E2 signaling and ERα signal at 24 h of E2 in WT-AR or DBM-AR overexpression on the TFF1 gene locus. **E.** Boxplot showing FOXA1 loss quantified by log2FC. FOXA1 (E2+DHT/E2) on overlapping and non-overlapping peaks. The y-axis shows fold change, and the x-axis denotes peak categories. **F.** Boxplot showing FOXA1 gain quantified by log2FC. FOXA1 (DBM-AR/WT-AR) on overlapping and non-overlapping peaks. The y-axis shows fold change, and the x-axis denotes the category of peaks. **G.** Boxplot showing ERα gain quantified by log2FC. ERα (DBM-AR/WT-AR) on overlapping and non-overlapping peaks. The y-axis shows fold change, and the x-axis denotes the category of peaks. **H.** Boxplot of AR signal at 24 h on ERα peaks showing increase, no change, and decrease based on log2FC. FOXA1 (E2+DHT/E2). The y-axis shows the AR levels at 24 h, and the x-axis denotes the different conditions. **I.** Boxplot of AR signal at 24 h on ERα peaks showing increase, no change, and decrease based on log2FC. FOXA1 (DBM-AR/WT-AR). The y-axis shows the AR levels at 24 h, and the x-axis denotes the different conditions. **J.** Boxplot of AR signal at 24 h on ERα peaks showing increase, no change, and decrease based on log2FC. ERα (DBM-AR/WT-AR). The y-axis shows the AR levels at 24 h, and the x-axis denotes the different conditions. **K.** Western blot of MCF7 cell lysates showing ERα and AR protein levels at 24 h of E2, following overexpression of WT-AR, A573D DNA-binding mutant (DBM) AR, or empty vector. The blot confirms overexpression of both WT-AR and A573D-AR, while total ERα protein levels remain unchanged upon AR overexpression. GAPDH was used as a loading control.

**Table S1:**
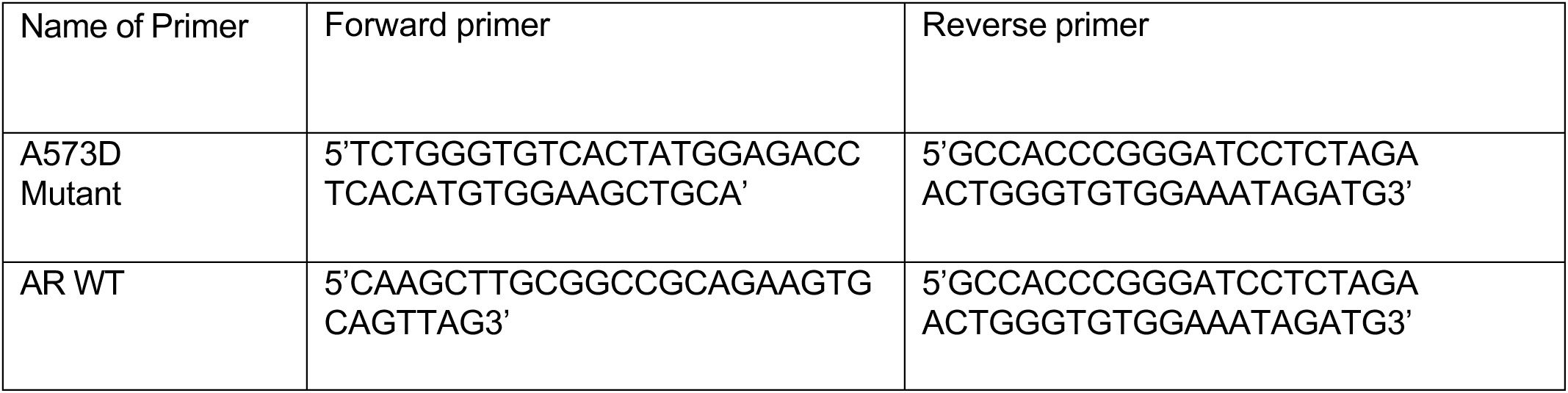
Primers used for cloning AR WT and A573D DNA-binding mutant.

**Table S2:**
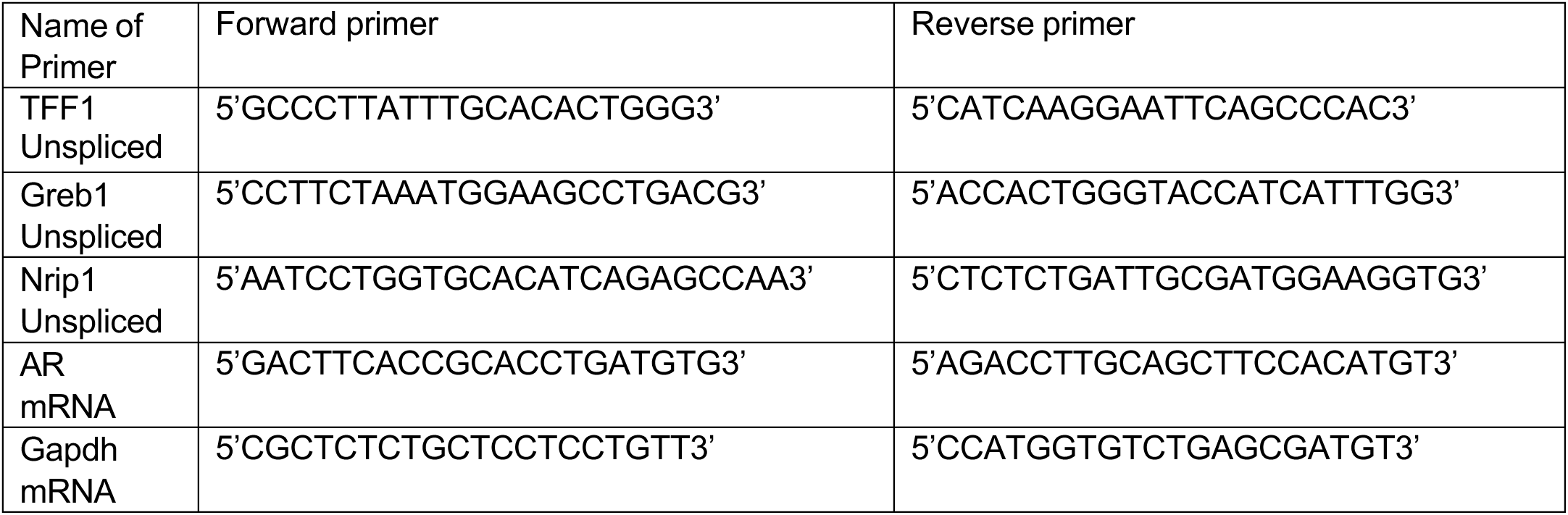
Primers used for evaluating ERα-mediated gene expression.

**Table S3:**
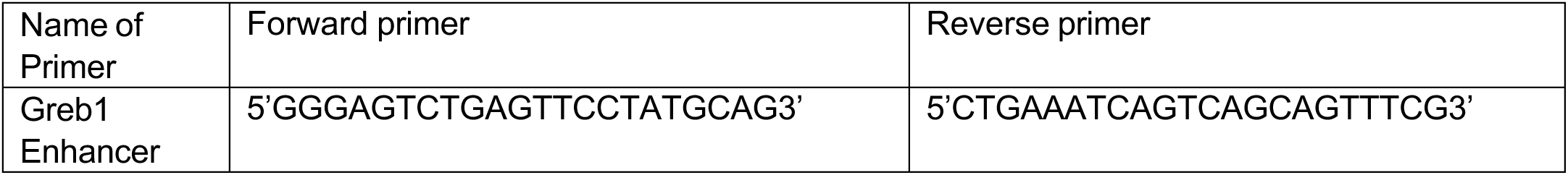
Primers used for evaluating anti-Flag ChIP-qPCR.

**Table S4:** Accession Number for GEO Datasets used in the study.

| Accession Number | Experiment | Reference |
| --- | --- | --- |
| GSM2467220 | MCF7_NoTreat_ERalpha | Dzida T. et al., 2017 |
| GSM2467222 | MCF7_E2_10min_ERalpha | Dzida T. et al., 2017 |
| GSM2467223 | MCF7_E2_20min_ERalpha | Dzida T. et al., 2017 |
| GSM2467224 | MCF7_E2_40min_ERalpha | Dzida T. et al., 2017 |
| GSM2467225 | MCF7_E2_80min_ERalpha | Dzida T. et al., 2017 |
| GSM2467226 | MCF7_E2_160min_ERalpha | Dzida T. et al., 2017 |
| GSM2467229 | MCF7_E2_1280_min_ERalpha | Dzida T. et al., 2017 |
| GSM1115992 | H3K27ac_ChIPSeq_E2 | Li W, Notani D. et al., 2013 |
| GSM1470000 | Med1_ChIPseq_E2_Exp9 | Liu Z. et al., 2014 |
| GSM1115995 | GROSeq_E2_repeat1 | Li W, Notani D. et al., 2013 |
| GSM1115996 | GROSeq_E2_repeat2 | Li W, Notani D. et al., 2013 |
| GSM678535 | GRO-seq_Vehicle_rep1 | Hah N. et al., 2011 |
| GSM678536 | GRO-seq_Vehicle_rep2 | Hah N. et al., 2011 |
| GSM678537 | GRO-seq_E2_10m_rep1 | Hah N. et al., 2011 |
| GSM678538 | GRO-seq_E2_10m_rep2 | Hah N. et al., 2011 |
| GSM678539 | GRO-seq_E2_40m_rep1 | Hah N. et al., 2011 |
| GSM678540 | GRO-seq_E2_40m_rep2 | Hah N. et al., 2011 |
| GSM678541 | GRO-seq_E2_160m_rep1 | Hah N. et al., 2011 |
| GSM678542 | GRO-seq_E2_160m_rep2 | Hah N. et al., 2011 |
| GSM1470014 | p300_ChIPseq_E2_Exp14 | Liu Z. et al., 2014 |
| GSM365930 | RNAPII E2 | Welboren et al. 2009 |
| GSM3728285 | AR Chip seq T47D | Hickey et al. 2021 |
| GSM3728287 | ER Chip seq T47D | Hickey et al. 2021 |
| GSM1589474 | H3K27ac seq T47D | Wang et al. 2015 |
| GSE341790 | ATAC_seq_E2_1hr_rep1 | This Study |
| GSE341790 | ATAC_seq_E2_24hr_rep1 | This Study |
| GSE341790 | ATAC_seq_E2_24hr_scr_rep1 | This Study |
| GSE341790 | ATAC_seq_E2_24hr_arkd_rep1 | This Study |
| GSE341790 | CHIP_AR_E2_1hr_rep1 | This Study |
| GSE341790 | CHIP_AR_E2_1hr_rep2 | This Study |
| GSE341790 | CHIP_AR_E2_24hr_rep1 | This Study |
| GSE341790 | CHIP_AR_E2_24hr_rep2 | This Study |
| GSE341790 | CHIP_ER_E2_1hr_rep1 | This Study |
| GSE341790 | CHIP_ER_E2_24hr_rep1 | This Study |
| GSE341790 | CHIP_ER_E2_1hr_rep2 | This Study |
| GSE341790 | CHIP_ER_E2_24hr_rep2 | This Study |
| GSE341790 | CHIP_ER_E2_1hr_resig_arkd_rep1 | This Study |
| GSE341790 | CHIP_ER_E2_1hr_resig_arkd_rep2 | This Study |
| GSE341790 | CHIP_ER_E2_1hr_resig_scr_rep1 | This Study |
| GSE341790 | CHIP_ER_E2_1hr_resig_scr_rep2 | This Study |
| GSE341790 | CHIP_ER_E2_24hr_AR573D_OE_rep1 | This Study |
| GSE341790 | CHIP_ER_E2_24hr_ARWT_OE_rep1 | This Study |
| GSE341790 | CHIP_FOXA1_E2_24hr_AR573D_OE_rep1 | This Study |
| GSE341790 | CHIP_FOXA1_E2_24hr_AR573D_OE_rep2 | This Study |
| GSE341790 | CHIP_FOXA1_E2_24hr_ARWT_OE_rep1 | This Study |
| GSE341790 | CHIP_FOXA1_E2_24hr_ARWT_OE_rep2 | This Study |
| GSE341790 | CHIP_FOXA1_E2_24hr_SCR_rep1 | This Study |
| GSE341790 | CHIP_FOXA1_E2_24hr_SCR_rep2 | This Study |
| GSE341790 | CHIP_FOXA1_E2_24hr_ARKD_rep1 | This Study |
| GSE341790 | CHIP_FOXA1_E2_24hr_ARKD_rep2 | This Study |
| GSE341790 | CutnRun_FOXA1_1hr_E2_rep1 | This Study |
| GSE341790 | CutnRun_FOXA1_1hr_E2_rep2 | This Study |
| GSE341790 | CutnRun_FOXA1_4hr_E2_rep1 | This Study |
| GSE341790 | CutnRun_FOXA1_4hr_E2_plus_DHT_rep1 | This Study |
| GSE341790 | CutnRun_FOXA1_4hr_E2_rep2 | This Study |
| GSE341790 | CutnRun_FOXA1_4hr_E2_plus_DHT_rep2 | This Study |
| GSE341790 | CutnRun_FOXA1_24hr_E2_rep1 | This Study |
| GSE341790 | CutnRun_FOXA1_24hr_E2_rep2 | This Study |
| GSE341790 | CutnRun_IgG_rep1 | This Study |
| GSE341790 | CutnRun_IgG_rep2 | This Study |

